# Single-Cell and Spatial Multi-omics Reveal Interferon Signaling in the Pathogenesis of Perianal Fistulizing Crohn’s Disease

**DOI:** 10.1101/2024.11.08.620717

**Authors:** Siyan Cao, Khai M. Nguyen, Kaiming Ma, Xin Yao, Ta-Chiang Liu, Malek Ayoub, Jalpa Devi, Yizhou Liu, Radhika Smith, Matthew Silviera, Steven R. Hunt, Paul E. Wise, Matthew G. Mutch, Sean C. Glasgow, William C. Chapman, Michelle Cowan, Mathew A. Ciorba, Marco Colonna, Parakkal Deepak

## Abstract

**Background & Aims:** Perianal fistulizing Crohn’s disease (PCD) is a common and debilitating complication with elusive pathophysiology. We examined mucosal cells from patients with PCD and related conditions using a multi-omics approach.

**Methods:** We recruited patients with PCD (n = 24), CD without perianal disease (NPCD, n = 10), and idiopathic perianal fistulas (IPF, n = 29). Biopsies were taken from fistula tracts, fistula opening, and rectal mucosa. Single-cell RNA-sequencing (scRNA-seq), mass cytometry (CyTOF), spatial transcriptomics (ST), immunohistochemistry, and integrated analysis were performed.

**Results:** ScRNA-seq, CyTOF, and ST unraveled immune and non-immune cell compartments in PCD and IPF fistula tracts. PCD fistulas showed hyperactivated pathogenic pathways including interferon (IFN)G response and TNF signaling in myeloid and stromal cells. Intestinal cells from PCD patients also expressed greater levels of IFNG-responsive and EMT genes compared to NPCD patients. Furthermore, both fistula tracts and ileal mucosa from PCD patients harbored expanded IFNG+ pathogenic Th17 cells, which expressed elevated inflammatory mediators. CyTOF also identified skewed immune cell phenotypes in the fistula tracts, fistula opening, and rectum in PCD patients including expanded Th17 cells, increased pathogenic myeloid cells, and altered T cell exhaustion markers. Further analysis also revealed cellular modules associated with anti-TNF therapy in PCD patients.

**Conclusion:** Multi-omics analysis revealed immune, stromal, and epithelial cell landscapes of PCD, which highlight the pathogenic role of hyperactivated IFNG signaling in both fistula tracts and luminal mucosa. This study identified IFNG as a potential therapeutic target for PCD.

## Introduction

Perianal fistulas occur in approximately up to 40% of patients with Crohn’s disease (CD) and are associated with high morbidity and an impaired quality of life^1^. The management of perianal fistulizing CD (PCD) is a major clinical challenge due to its refractoriness to medical treatment despite recent advancement in targeted therapies, where only one-third of the patients achieve remission at one year^1^ ^2^. The underlying etiology of PCD remains poorly understood, which hinders the development of therapies that specifically target this debilitating complication^3^. An earlier study showed that PCD fistulas exhibited altered populations of CD45RO+ T cells, CD20+ B cell, and CD68+ macrophages in histopathology^4^. An increased IL-12p70 was found at the internal opening of PCD fistulas compared to those with idiopathic perianal fistulas (IPF)^5^. Recently, Levantovsky et al. showed enrichment of myeloid cells and CHI3L1^hi^ fibroblasts and altered myeloid-stromal crosstalk in fistula tracts vs. colorectal tissues from three PCD patients using multimodal single-cell analyses^6^. Using flow cytometry and bulk RNA-seq, Constable et al. identified increased CD161+ CD4+ T cells and CD161+ CD4-CD8-invariant NKT cells that produced IL-22 and IL-13 in PCD fistula tissues, as well as elevated expression of genes associated with epithelial-to-mesenchymal transition (EMT) and tissue remodeling in PCD^7^.

Despite those findings, a detailed picture of the mucosal cell populations in the fistula tracts and nearby tissues in PCD versus related conditions has been lacking. In this study, we used a multi-omics approach including single-cell RNA-sequencing (scRNA-seq), mass cytometry (CyTOF), and spatial transcriptomics (ST) to unravel mucosal cell response in an extensive cohort of patients with PCD, CD without perianal disease (NPCD), or idiopathic perianal fistula or cryptoglandular fistulas (IPF). PCD fistulas showed hyperactivated pathogenic signals including interferon (IFN)G and TNF responses. Ileal and rectal cells from PCD patients also expressed increased IFN-γ-responsive and EMT genes compared to NPCD patients. Furthermore, we identified expanded IFNG+ pathogenic Th17 cells in both fistula tracts and ileal mucosa from PCD patients. Altered frequencies and inflammatory phenotypes of immune cells in the fistula tracts, external fistula opening, and rectum in PCD patients were uncovered by CyTOF. Moreover, molecular modules associated with anti-TNF therapy were identified in PCD samples. Altogether, this study identified IFN-γ as a potential therapeutic target for PCD.

## Materials and Methods

### Patient Recruitment and Sample Collection

We recruited three groups of patients with a history of: 1) PCD confirmed by magnetic resonance imaging (MRI) of the pelvis and/or examination under anesthesia (EUA) by a colorectal surgeon (n = 24); 2) NPCD (n = 29); 3) IPF without a history of inflammatory bowel disease, diagnosed on EUA by a colorectal surgeon (n = 10; **Supplementary Table 1**). Individuals with a prior diagnosis of secondary perianal fistulas related to infection or malignancy, active infection or malignancy involving the anorectal area, and/or contraindication to biopsies (e.g. bleeding risks, immunodeficiency, or other contraindications deemed by the surgeons who perform EUA) were excluded. Patients who were less than 18 years of age, pregnant, or unable to provide informed consent were also excluded. Three biopsies each were taken using endoscopic biopsy forceps from the fistula tracts, external opening of the fistula, and/or nearby rectal mucosa (near internal fistula opening) from patients in PCD and IPF groups during routine EUA. Random rectal biopsies were taken from patients with NPCD during a routine colonoscopy. For each location sampled, biopsies were also collected for routine pathology assessment by two GI pathologists. Samples were cryopreserved within one hour of collection as previously described^8^. Formalin-fixed paraffin-embedded tissue blocks of fistula tracts from 6 patients with PCD were obtained from the Division of Anatomic & Molecular Pathology at Washington University School of Medicine.

### CyTOF

Cells were washed with Cy-FACS buffer (CyPBS, Rockland, MB-008; 0.1% BSA, Sigma, A3059; 0.02% Sodium Azide, Sigma, 71289, 2mM EDTA, Hoefer, GR123-100) and stained with antibodies on ice for an hour. After two washes cells were stained with cisplatin (Enzo Life Sciences, NC0503617) for one min to label dead cells. They were then washed again twice, fixed in 4% PFA (Electron Microscopy Sciences, 15710) for 15 min, spun down and stained with barcodes for 30 min. The cells were then re-suspended in Intercalator-Ir125 (Fluidigm, 201192 A) overnight. Cells were washed, counted and analyzed by certified staff using CyTOF2/Helios instrument (Fluidigm) at the Immunomonitoring Laboratory at Washington University as previously described^8, 9^. Data was analyzed using Cytobank, and significant differences in cell frequencies were determined by one-way ANOVA. Additional details on data analysis and gating are included in **Supplemental Methods.**

### Single cell and spatial transcriptomics analyses

Samples were processed using Cellranger (v7.0.0) by the Genome Technology Access Center at Washington University. Cellranger output was loaded and analyzed in R using Seurat (v4.4.0). ST data was processed using SpaceRanger (v3.0.1) and analyzed using Seurat. Cell type deconvolution was performed using CARD (v1.1)^10^, and ligand-receptor signaling was inferred using NICHES (v1.1)^11^. Details about library preparation and data analyses are included in **Supplemental Methods**.

## Results

### Single-cell transcriptomics resolves cellular heterogeneity in perianal fistulas

Fistula tract biopsies of 9 with PCD and 6 individuals with IPF were analyzed using scRNA-seq. Following strict quality control, a total of 56,560 high-quality cells were retained for subsequent analyses. The major cell compartments were resolved based on canonical markers, including T cells (CD3D+) and innate lymphoid cells (ILCs), B cells (MS4A1+), plasma cells (SDC1+), monocytes/macrophages/dendritic cells (ITGAX+), stromal cells (COL1A1), and endothelial cells (VWF+) (**Figure 1A and Supplementary Figure 1**).

**Figure 1.**
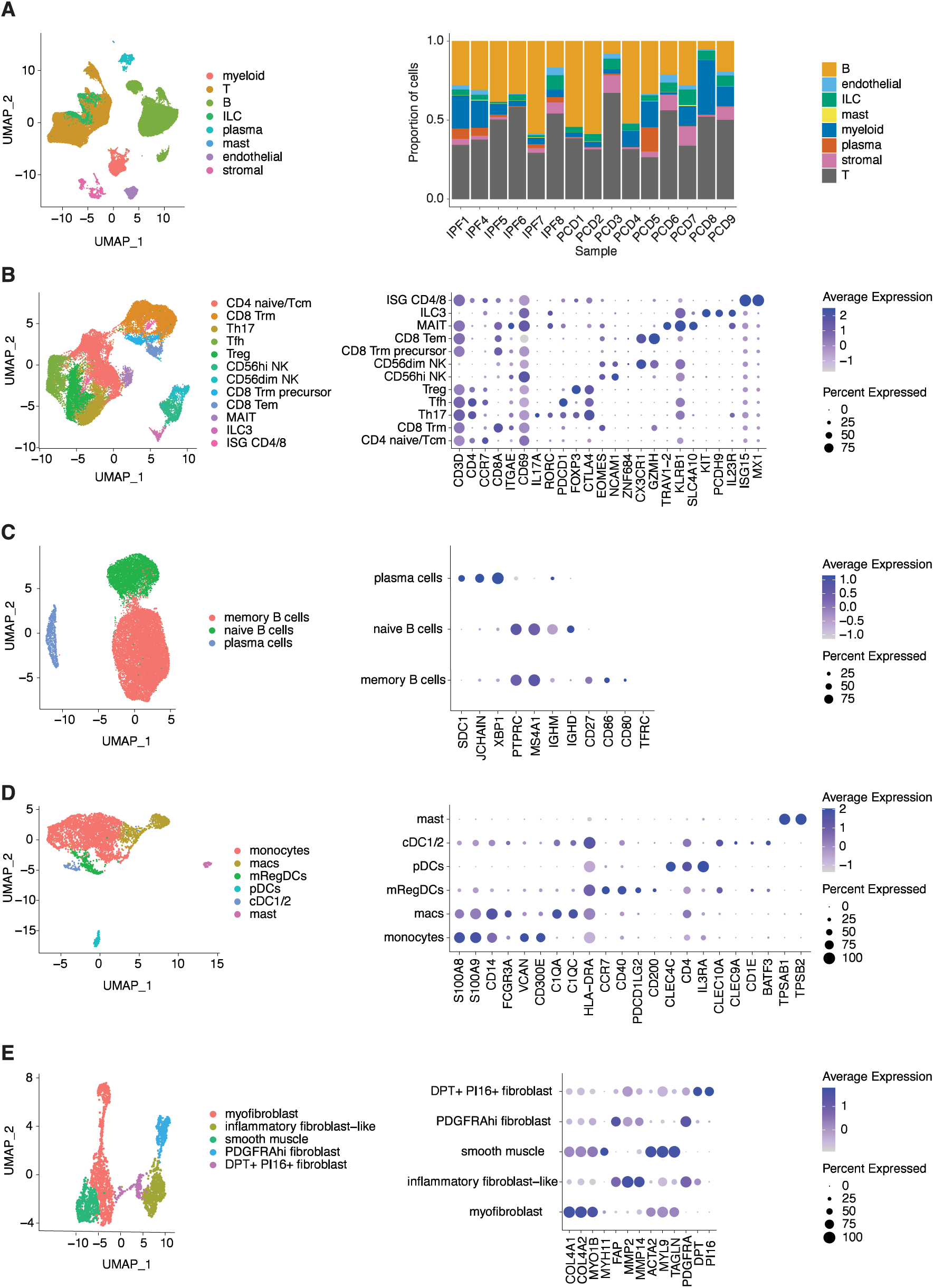
Cellular heterogeneity of fistula tracts revealed by scRNA-seq. (A) Left: UMAP plot of major cell compartments, right: Cell type composition across samples. (B-E) Left: UMAP plots, right: marker genes used to identify subsets of T cells and ILCs (B), B cells and plasma cells (C), myeloid cells (D), and stromal cells (E).

We separately re-clustered T cells and ILCs (T-ILC), B cells and plasma cells, myeloid cells, and stromal cells (**Figure 1B-E; Supplementary Table 2**). T and innate lymphoid cell (ILC) clusters consisted of naïve CD4 (CD4+, CCR7+), regulatory T cells (Treg; FOXP3+ CTLA4+), T-helper 17 cells (Th17; CD4+ IL17A+ RORC+), follicular helper T cells (Tfh; CD4+ PDCD1+), mucosal-associated invariant T cells (MAIT; TRAV1-2+ KLRB1+ SLC4A10+), effector memory CD8 T cells (CD8 Tem; CD8+ CX3CR1+ GZMH+), resident memory CD8 T cells (Trm; CD8+ ITGAE+ CD69+), Trm precursor cells^12^ (CD8+ ZNF683+), group 3 ILCs (ILC3; CD3D-KIT+ PCDH9+ IL23R+ RORC+), CD56^hi^ NK cells (CD3D-EOMES+ NCAM1^high^), and CD56^low^ NK cells (CD3D-EOMES+ NCAM1^low^) (**Figure 1B**). B cells included naïve B cells (IGHM+ IGHD+) and memory B cells (CD27+ CD86+ CD80+ TFRC+) **(Figure 1C)**. Myeloid cells included monocytes (CD14+ FCGR3A+ S100A8+ VCAN+), macrophages (C1QA+ C1QC+ HLA-DRA+), conventional dendritic cells (cDC; CLEC10A+ CLEC9A+ CD1E+ BATF3+), plasmacytoid dendritic cells (pDC; CLEC4C+ CD4+ IL3RA+), mature DC enriched in immunoregulatory molecules^13^ (mRegDCs; CCR7+ CD40+ PDCD1LG2+ CD200+), and mast cells (KIT+ TPSB2+ TPSAB1+) (**Figure 1D**). Stromal cells included smooth muscle cells (ACTA2+ MYL9+ TAGLN+), inflammatory fibroblast-like cells (FAP+ MMP2+ MMP14+), PDGFRA^hi^ fibroblasts (PDGFRA^hi^), DPT+ PI16+ fibroblasts (DPT+ PI16+), and myofibroblasts (COL4A1+ COL4A2+ MYO1B+ MYH11+) (**Figure 1E**).

### PCD is marked by enhanced IFNG and TNF responses

To identify pathways that define PCD fistulas at a global level, we performed single-cell pathway analysis (SCPA)^14^ to compare PCD to IPF. IFN-γ and TNF-α signaling were among pathways increased activity in PCD (**Figure 2A**). In contrast, type I IFN pathway did not exhibit enhanced activity (**Supplementary Figure 2A**). To determine whether activation of IFN-γ pathway was fistula-specific in PCD, we repeated SCPA on published scRNA-seq datasets of rectum^15^ (active PCD vs. inactive/healed PCD; n = 6 per group) as well as colon and terminal ileum^16^ (PCD vs. NPCD; n = 8 for PCD, n = 39 for NPCD). Elevated IFN-γ and TNF-α pathway activity was seen in all three gastrointestinal tract locations in PCD (**Figure 2B**). We validated upregulation of IFN-γ pathway by showing increased IFN-γ and phosphorylated STAT1 (downstream of IFN-γ) in PCD tissues using IHC (**Figure 2C**). Taken together, PCD is characterized by heightened IFN-γ and TNF-α responses in both fistula tracts and intestinal mucosa.

**Figure 2.**
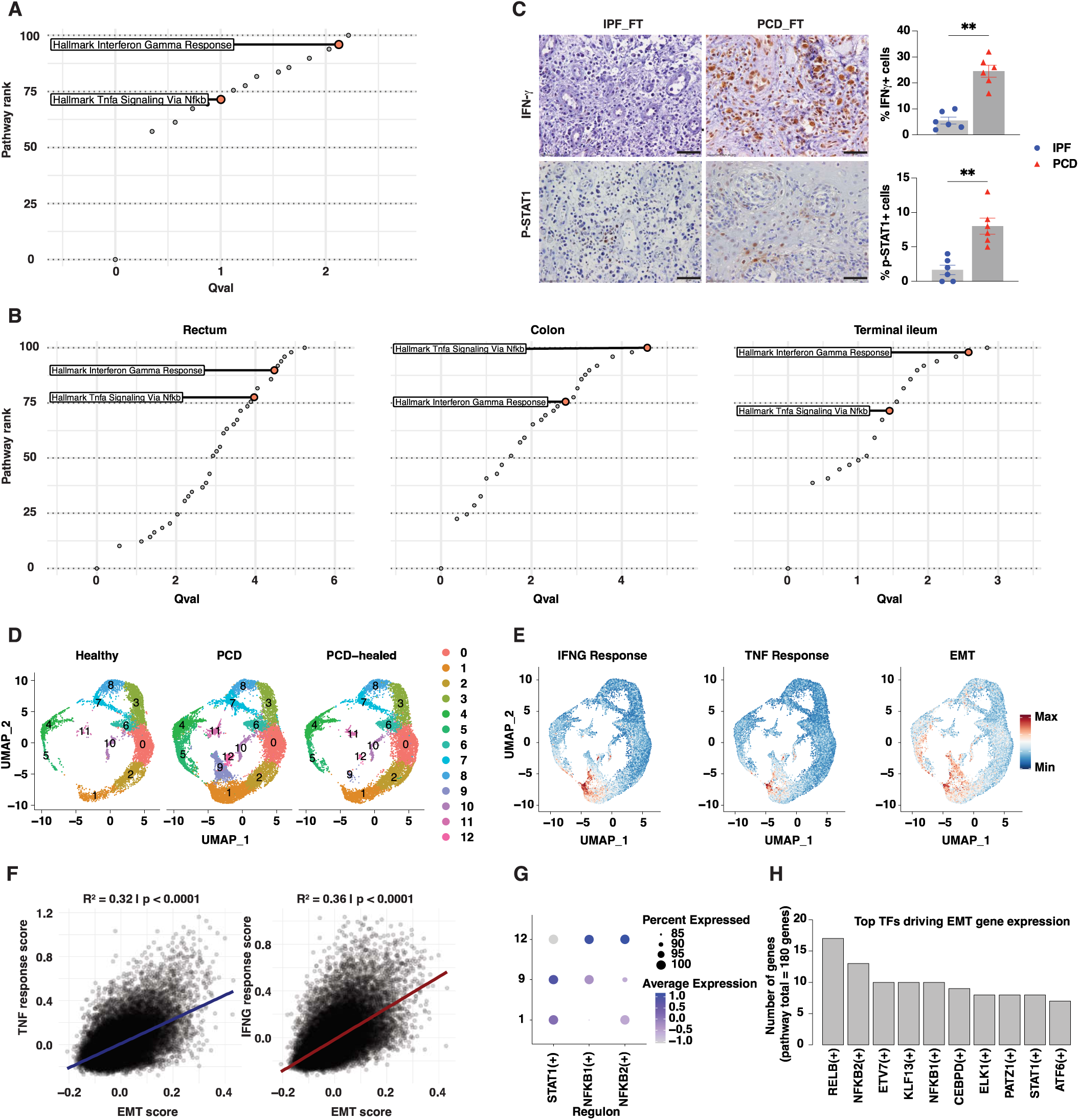
IFN-γ signaling is a distinguishing feature of PCD. (A) Pathway rank plot of top upregulated pathways in PCD fistulas compared to IPF fistulas. (B) Pathway rank plot of top upregulated pathways in rectum (active PCD vs. healed/inactive PCD) as well as colon and terminal ileum (PCD vs. NPCD). Complete SCPA output as in (A) and (B) can be found in Supplementary Table 3. (C) Representative IHC image and quantification of IFN-γ and p-STAT1 staining of IPF and PCD fistula tracts (FT). (D) UMAP plots comparing rectal epithelial cell clusters between healthy controls, PCD, and PCD-healed. (E) UMAP plots of single-cell module scores for IFNG response, TNF response, and EMT pathways in PCD rectal epithelial cells. (F) Correlation between TNF (left) or IFNG (right) response and EMT scores. (G) Dot plot indicating transcription factory activity of STAT1, NFKB1, and NFKB2 in rectal epithelial cell clusters 1, 9, and 12 as in (D). (H) Bar plot of top transcription factors driving EMT gene expression in rectal epithelial cells. ** p<0.01.

EMT has been implicated in the pathogenesis of PCD^3, 17, 18^. We hypothesized that inflammatory signaling by TNF-α and/or IFN-γ in PCD induce EMT gene expression. To understand their relationships, we first subsetted and re-clustered epithelial cells from the rectum of PCD patients^15^ (**Supplementary Figure 2B; Supplementary Methods**). Compared to uninflamed samples, inflamed epithelium (proctitis) was enriched in cluster 9 (C9) that consisted of cells expressing IFN-stimulated genes, as well as C12 marked by multiple cytokine genes (**Figure 2D and Supplementary Figure 2C**). Interestingly, C9 and C12 clustered closely to C1, which harbored colonocytes expressing both subunits of IFN-γ receptor (IFNGR1 and IFNGR2) as well as TNF-α receptor (TNFRSF1A) (**Supplementary Figure 2C**). Single-cell scores for responses to IFN-γ, TNF-α, and EMT were all elevated in C9 and C12 compared to the remaining clusters (**Figure 2E**), and IFN-γ, and TNF-α response scores were significantly correlated with EMT score (**Figure 2F)**. SCENIC analysis of transcriptional networks^19^ revealed elevated transcription driven by STAT1 in C9 (**Figure 2G**). Similarly, NFKB1 and NFKB2, which are downstream of both IFN-γ and TNF-α signaling^20–22^, were increased in C12. Cross-comparison of STAT1, NFKB1, and NFKB2 target genes with the GO EMT gene set revealed these transcription factors as top drivers of EMT gene expression (**Figure 2H**). Thus, our analysis suggests direct regulation of EMT by IFN-γ and TNF-α signaling in PCD.

### Pathogenic Th17 cells and myeloid cells contribute to IFNG signaling in PCD

Ligand-receptor signaling analysis was performed using NICHES^11^ to resolve cell-cell interactions in an unbiased manner. Top ligand-receptor pairs enriched in PCD included IFNG-IFNGR2, GZMB-PGRMC1, IL2-IL2RG, MMP9-CD44, VIM-CD44, CCL4-CCR5, CCL5-CCR5, among others (**Figure 3A; Supplementary Table 4**). IFNG, GZMB, and IL2 indicate paracrine signaling of immune cells; MMP9 and VIM highlight stromal remodeling and EMT; and the chemokines CCL4 and CCL5 suggest recruitment of immune cells to the fistula tracts. While TNF-α is known to be pathogenic in PCD^23^, TNF signaling was present in both IPF and PCD (**Supplementary Figure 3A**). A recent study reported upregulation of IL22 and IL13 in PCD relative to IPF^7^; indeed, we found an increased cell-cell interactions for both IL22 and IL13 signaling in PCD (**Supplementary Figure 3B**). We further analyzed the IFNG pathway to determine cell types underlying elevated IFN-γ signaling in PCD. IFNG senders comprised of heterogenous cell populations; however, only Th17 and regulatory T cells (Tregs) were significantly enriched in PCD (**Figure 3B**). IFN-γ-producing Th17 cells represent a pathogenic antigen-induced state^24–26^, while IFN-γ expression in Tregs can be induced by IFN-γ itself^27, 28^. Thus, these data suggest that pathogenic Th17 cells (pTh17) underlie the hyperactivated IFNG response in PCD.

**Figure 3.**
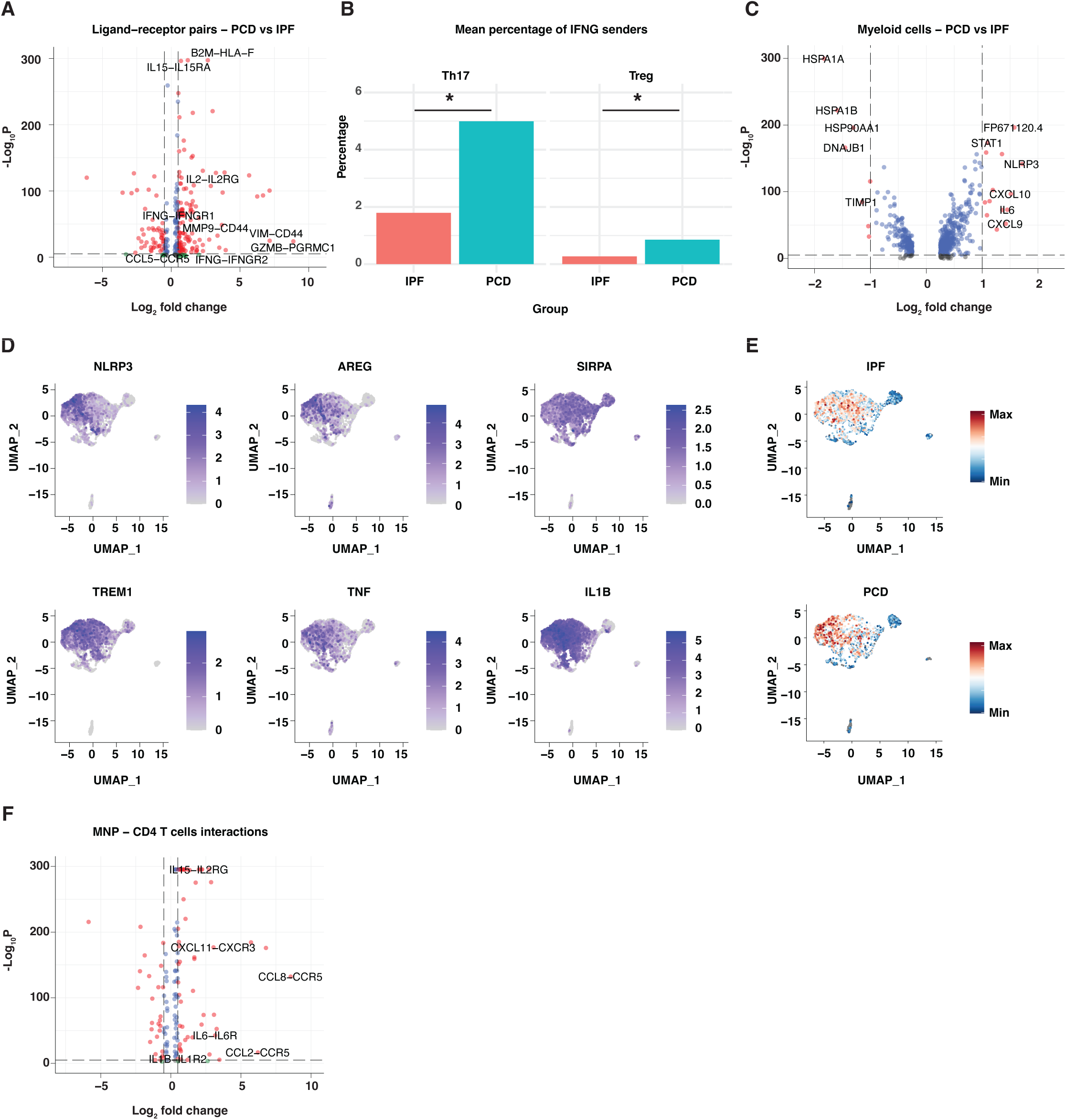
Pathogenic myeloid cell-Th17 cell interaction underlies IFN-γ response in PCD fistulas. (A) Volcano plot highlighting select upregulated ligand-receptor signaling pairs. (B) Bar plot indicating IFNG senders significantly enriched in PCD. (C) Volcano plot of differentially expressed myeloid cell genes between PCD and IPF. (D) UMAP plots of NLRP3, AREG, SIRPA (CD172a), TREM1, TNF, and IL1B expression in myeloid cells. (E) UMAP plots of single-cell module scores for LPS signaling in IPF and PCD myeloid cells. (F) Volcano plot showing top enriched ligand-receptor signaling pairs between mononuclear phagocytes (MNPs) and CD4 T cells in PCD. * p<0.05

As pTh17 cells are likely a product of cytokine and antigen stimulation, we next analyzed the antigen presenting cells (APCs) in our dataset. While there were few differentially expressed genes (DEGs) in B cells (**Supplementary Figure 3C, D**), myeloid cells expressed increased NLRP3, AREG, and IFN-γ-responsive chemokines CXCL9 and CXCL10 (**Figure 3C**). Additionally, NLRP3+ AREG+ cells also expressed TREM1 and SIRPA (encoding CD172a) (**Figure 3D**). We next calculated single-cell scores for signaling by LPS, a membrane component of gram-negative bacteria and toll-like receptor (TLR)4 ligand, revealing an upregulation of this pathway in NLRP3+ AREG+ cells and the entire myeloid compartment in PCD compared to IPF (**Figure 3E**). In human monocytes, LPS leads to alternative/non-canonical activation of NLRP3 inflammasome independently of potasium efflux and pyroptosis^29^. Notably, NLRP3 is also transcriptionally induced via TLR4 activation^29, 30^. Ligand-receptor analysis showed activation of multiple APC-derived signals towards naïve CD4 and Th17 cells (**Figure 3F; Supplementary Table 4**), including chemokines (CCL8, CCL2, CXCL10, CXCL11, CXCL9) and cytokines (IL6, IL15, IL1B). These chemokines function to recruit T cells to site of inflammation^31, 32^, with CXCL9/10/11 being crucial in IFN-γ-skewed T cell responses^33^. IL6 and IL1B are key signals in Th17 differentiation from naïve T cells^24^, while IL15 has been shown to promote Th17 responses by inducing RORγt^34^. Altogether, these data highlight the role of myeloid cells in the recruitment and activation of pathogenic T cells including Th17 cells, and support LPS as a potential trigger of PCD.

### CyTOF uncovers pathogenic immune responses in PCD fistula tracts and surrounding tissues

To validate the immunophenotype of PCD at the protein level, we analyzed mucosal immune populations from multiple locations in patients with PCD, NPCD, or IPF using CyTOF **(Supplementary Figure 4A)**. In the fistula tract, CD4 T cells expressing the activating markers CD137 and CD56 were elevated in PCD relative to IPF (**Figure 4A**). CD56+ CD4 T cells represent a morphologically distinct population with a mature, non-proliferative phenotype and enhanced production of IFN-γ and TNF-α^35^, while CD137 is expressed on activated T cells and increased in lamina propria cells in CD tissues^36^. Circulating Th17 cells were also elevated (**Figure 4B**); increased recruitment of these cells to the fistula site is likely due to increased chemokine signaling, as shown in our ligand-receptor analysis. Similarly, we found an increase in CD172a/b+ TREM1+ CD14+ mononuclear phagocytes (MNPs; **Figure 4C**), which has been reported to be pathogenic in luminal CD^37^. CD45+ cells, CD19+ B cells, and CD45RO+ memory T cells were also elevated in PCD fistulas (**Supplementary Figure 4B**), consistent with a prior histological study^38^. Additionally, exhaustion marker CD39 was increased while CD127 was decreased in both CD4 and CD8 T cells from PCD fistula samples (**Figure 4D**). This is important because T cell exhaustion has been linked to prolonged IFN-γ production in prior studies^39–41^. To understand the pathologic relationship between fistula tracts and surrounding tissues in PCD, we analyzed cells from the external perianal fistula opening of PCD and IPF patients. The frequencies of CD45RO+ T cells was higher in PCD external fistula opening vs. that of IPF (**Supplementary Figure 4C**), similar to the changes in fistula tracts. Expression of CD127 in CD4 and CD8 T cell subpopulations also trended decreasing in PCD external fistula opening (**Supplementary Figure 4D**). In contrast to the fistula tracts, the specimens from PCD external fistula opening exhibited increased TRM CD8 T cells compared to those from IPF (**Supplementary Figure 4E**). Pathogenic CD172a/b+ TREM1+ MNPs were also enriched in PCD external fistula opening (**Figure 4E**). Lastly, we characterized rectal immune cells from patients with PCD, NPCD, and IPF. Like the fistula tracts, PCD rectum exhibited higher CD39 level in both CD4 and CD8 T cells compared to those from NPCD and IPF (**Figure 4F**). Rectal CD172a/b+ TREM1+ MNPs were more abundant in PCD (**Figure 4G**), consistent with the findings in both fistula tracts and external fistula opening. IL-17-producing CD8 T cells (Tc17; marked by CD26 and CD161) were also enriched in PCD rectum relative to both IPF and NPCD (**Supplementary Figure 4F**). Taken together, our CyTOF data corroborated transcriptomic findings and revealed additional characteristics indicative of IFN-γ signaling in PCD fistula tracts, external fistula opening, and rectum.

**Figure 4.**
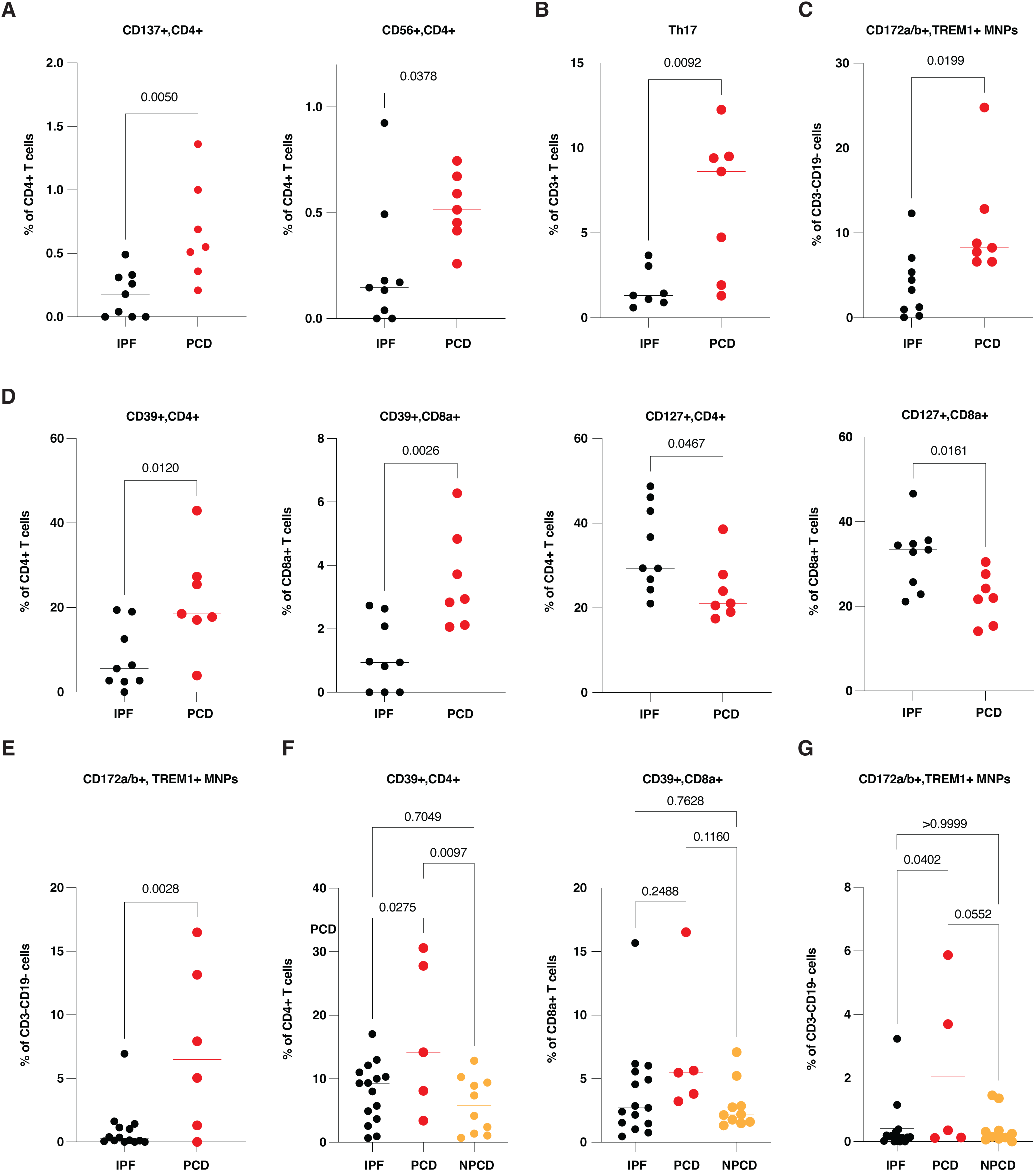
CyTOF revealed immunologic relationship between fistula tracts and surrounding tissues. Plots of CyTOF data comparing cell frequencies in fistula tracts (A-D), external fistula opening (E), and rectum (F, G): (A) CD137+ and CD56+ CD4 T cells. (B) Th17 cells. (C) CD172a/b+ TREM1+ MNPs. (D) CD127 and CD39 expression on CD4 and CD8 T cells. (E) CD172a/b+ TREM1+ MNPs. (F) CD39 expression on CD4 and CD8 T cells. (G) CD172a/b+ TREM1+ MNPs.

### Transcriptomic alterations in PCD fistulas following anti-TNF treatment

At present, anti-TNFs, especially infliximab, have the best evidence for the treatment of PCD^3, 23, 42^. However, the mechanisms by which they induce fistula healing remain unclear. To address this unmet need, we divided our scRNA-seq cohort into those on an anti-TNF agent or advanced therapy-naïve groups (n=4/group), based on their treatment at time of sampling. Weighted gene co-expression network analysis was performed on pseudobulked scRNA-seq data to identify gene modules associated with anti-TNF treatment^43^. In total, 32 gene modules were identified and two were significantly correlated with anti-TNF treatment (**Figure 5A; Supplementary Table 5**). The “lightpink1” module, which was downregulated with TNF antagonists, was comprised of multiple genes associated with T cells (CD3E, CD3D, TNFRSF9, LCK, CD247, IKZF2, TCF7, LAT, and TCR genes), myeloid cells (C1QA, C1QC, C1QB, IL6ST, AREG, CCL22, BATF3), and interferon response (IFITM2, IFITM1, IRF3) (**Figure 5B**). The “royalblue2” module, which was upregulated with anti-TNF use, consisted of genes encoding structural proteins (GJA9, OCLN, FSCN1, CLDN15, MKLN1, NEFL), proteins involved in cellular metabolism (SLC1A7, HCN3, GTDC1, OXSM, SFXN3, CASR, SLC25A16, CARNS1), and DNA replication, transcription and mRNA translation (SNHG3, LIN54, ZGRF1, TMA16, RPL26L1, HIST1H4I, ZSCAN9), which were likely linked to cell proliferation concomitant with fistula healing (**Figure 5C**). Notably, the “royalblue2” module also contained MAP3K21 (encoding MLK4), a negative regulator of TLR4 signaling^44^; WDR41, which protects against murine colitis via TLR signaling^45^; and CCL27, a key chemokine for immune homeostasis^46^. Thus, our data suggest that anti-TNF treatment leads to fistula healing by suppressing pathogenic T-cell and myeloid-cell signatures, while enhancing cell proliferation, mucosal barrier integrity, and immunoregulatory processes.

**Figure 5.**
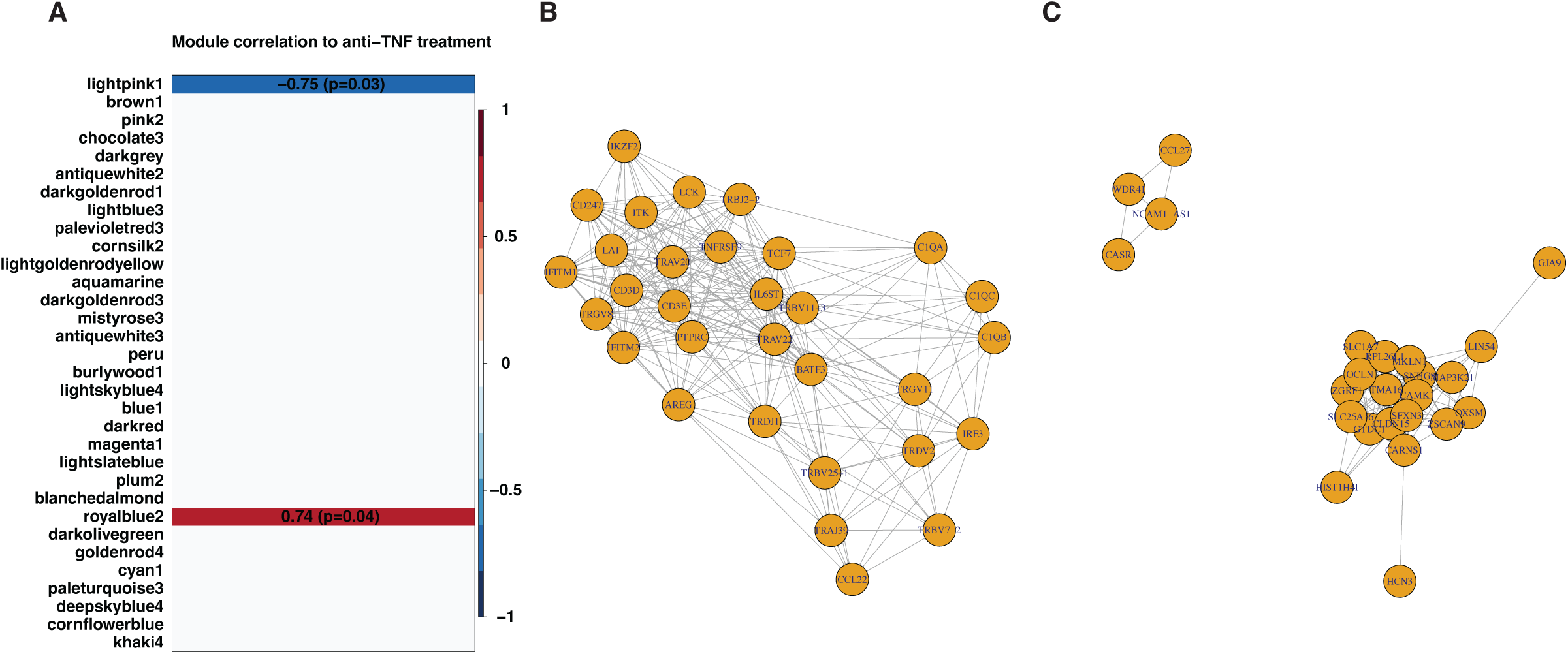
Gene expression modules in PCD fistula tracts correlated with anti-TNF therapy. (A) Co-expression gene modules identified by WGCNA. Significantly correlated modules are colored: red - positive correlation; blue - negative correlation. P-values denoted by number in parentheses. (B) Network of key lightpink1 module members associated with T and myeloid cell function. (C) Network of key royalblue2 module members associated with structural integrity, metabolism, proliferation, and immune homeostasis.

### Spatial transcriptomics resolves shared and unique microenvironments in IPF and PCD fistulas

To better understand the spatial transcriptomic dynamics in perianal fistula, we performed ST on FFPE blocks of IPF and PCD fistula tracts using 10X Visium (n=3 per group; **Supplementary Figure 5A**), which generated a total of 27,226 spots. We first combined the datasets and performed integrated dimensionality reduction and clustering to identify shared transcriptomic niches, which identified six spatially correlated clusters **(Figure 6A**, **Supplementary Figure 5B)**. The fistula tract lining was mostly made up of Cluster 1 (C1), which was marked by transcripts of keratins (KRT17, KRT16, KRT13, KRT1, KRT14; **Supplementary Figure 6A**). C1 was interspersed with C0, which was marked by immune cell genes including those for macrophages (APOE, LYZ), B cells and plasma cells (IGHM, IGHG1, JCHAIN, IGHA1), and T cells (IL7R, LTB)(**Supplementary Figure 6A**). Notably, the fistula tracts in sample IPF1 were devoid of C1 spots but instead included C5, which was marked by expression of neutrophil/inflammatory monocyte markers and cytokines (LYZ, CXCL8, SOD2, ITGAX, IL1RN, IL1B, SPP1, MMP9) as well as foci of inflammation adjacent to the fistula tract (**Supplementary Figure 6A**). This was perhaps due to lack of epithelialization in this fistula tract, a phenomenon reported in a prior histological study^38^. C2 in IPF1 and PCD1 was made up of connective tissue distant from the fistula tract (**Supplementary Figure 5**), and marked by expression of extracellular matrix proteins (COL1A1, COL1A2, SPARC, COL6A2, DCN, DPT; **Supplementary Figure 6A**). C3 formed the connective tissue layer adjacent to the fistula tract, which was marked by expression of myofibroblast genes (MYL9, TAGLN, MYH11, DES, ACTA2; **Supplementary Figure 6A**). C4 harbored mixed features of C0, C3, and C5 and was only present in IPF1. Altogether, these data indicate shared transcriptomic niches across PCD and IPF fistula tracts.

**Figure 6.**
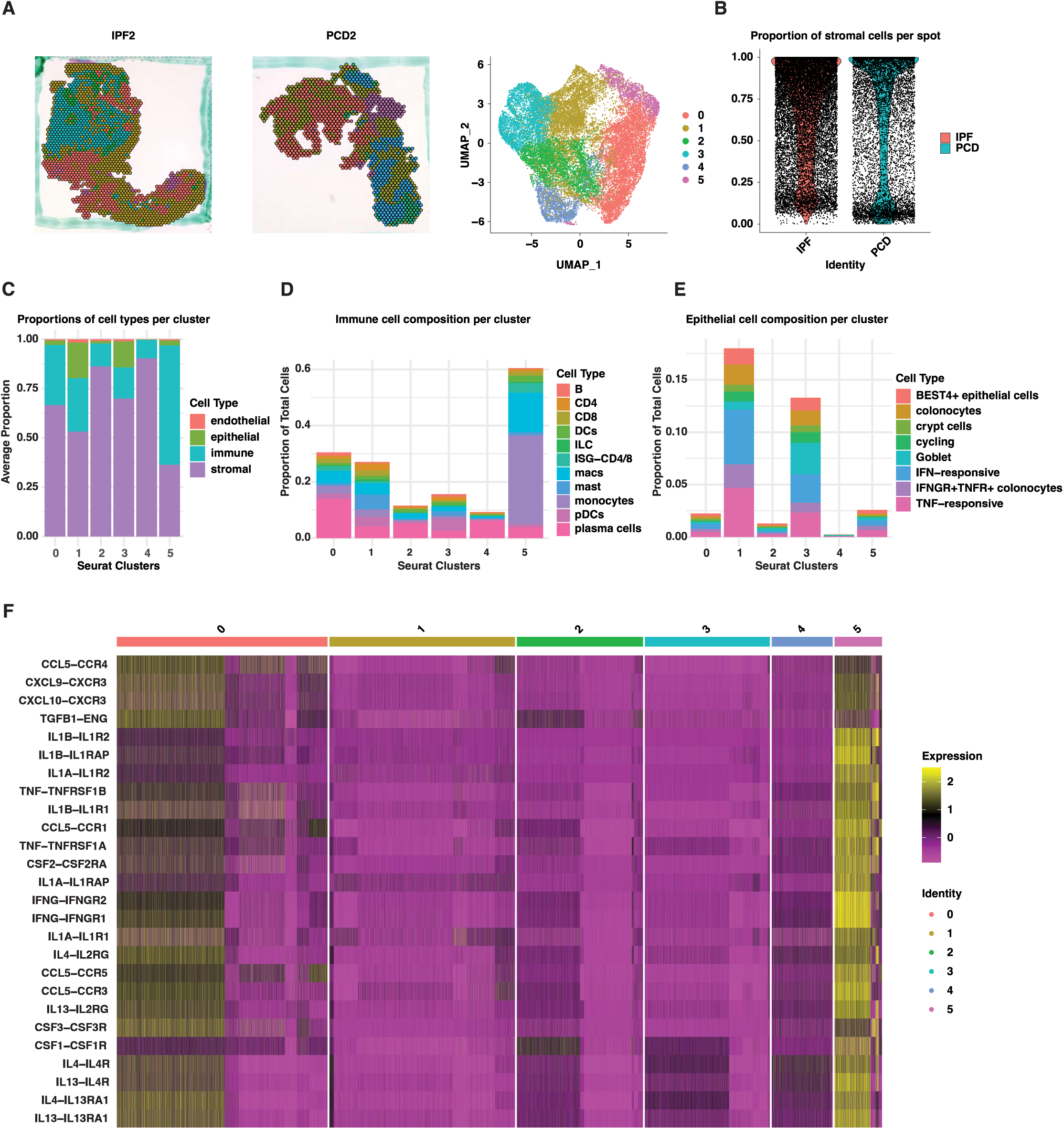
Spatial transcriptomics identified shared transcriptomic niches across fistula tracts. (A) Left: representative plots showing spatial organization of dots in each cluster in IPF and PCD fistulas; right: UMAP plot of spatial clusters. (B) Proportion of stromal cells per dot calculated by CARD. (C) Distribution of major cell compartments across spatial clusters. (D) Immune cell composition across spatial clusters. (E) Epithelial cell composition across spatial clusters. (F) Heatmap of cytokine ligand-receptor signaling across spatial clusters.

We next performed cell type deconvolution of the ST data using our single-cell data as reference^10^. Since our scRNA-seq dataset does not contain a significant epithelial cell population, probably due to non-epithelialized fistula tracts in that cohort, we integrated our dataset with a PCD rectal scRNA-seq dataset containing epithelial cells^15^ to ensure adequate deconvolution of fistula tract epithelium (**Supplemental Methods**). Most spots profiled in the fistulas contained significant proportions of fibroblasts (**Figure 6B**); C1 and C3 had highest proportions of epithelial cells; while C5 consisted of mostly immune cells (**Figure 6C**). We explored composition of immune cell subtypes within each cluster (**Figure 6D**). C0 comprised primarily of plasma cells, followed by macrophages and monocytes, while C1 harbored the highest proportion of mast cells. C5 was enriched in monocytes, macrophages and ISG-CD4/8 T cells, potentially resembling tertiary lymphoid structures that are aggregates of lymphoid and APCs commonly arising during chronic inflammation^47^. We also evaluated the composition of epithelial subtypes. Epithelial cells lining the fistula tract in C1 were primarily mapped to IFNGR+ TNFR+, IFNG- and TNF-responsive colonocytes, which corresponded to C1, C9 and C12, respectively (**Figure 6E**). These data further support the potential role of epithelial IFNG and TNF signaling in perianal fistulization.

Spatial ligand-receptor interactions were analyzed using NICHES^11^, which revealed distinct ligand-receptor signaling pairs across the spatial clusters (**Supplementary Figure 6B**). Filtering for cytokine ligand-receptor pairs identified two main foci of immune signaling (**Figure 6F**). C0, which was adjacent to the fistula tract, was marked by chemokines including CCL5, CXCL9, and CXCL10 signaling pathways. CCL5 is expressed by activated T cells and monocytes, and is broadly chemotactic for immune cells. CXCL9 and CXCL10 are notably induced by IFN-γ and amplify IFN-γ-producing T cells as well as chemoattraction of myeloid and lymphoid cells, respectively^48, 49^. Notably, C0 also overexpressed the WNT5A-ROR2 wound healing pathway, as well as the TGFB1-CXCR4 pathway which is implicated in EMT^50–52^. C5, on the other hand, was marked by > 100 cytokine ligand-receptor pairs including TNF, IFNG, IL1A/B, IL4, and IL13, as well as multiple chemokines. Interestingly, signaling by GM-CSF, M-CSF and G-CSF (encoded by CSF1, CSF2 and CSF3, respectively) were also enriched in C5; this may represent a supportive mechanism for myeloid cell survival following their infiltration from circulation. Collectively, these data suggest niche-specific recruitment and regulation of immune cells, which may contribute to maintenance of inflammatory microenvironment in perianal fistulas.

Finally, we examined spatial organization of PCD-specific features. Single-spot enrichment of signature gene modules for pTh17 and NLRP3+ AREG+ MNPs were calculated to infer spatial organization of these cell types. NLRP3+ AREG+ MNPs were more ubiquitously present, particularly at tract-adjacent spots, suggesting their interactions with microbial elements in the fistula tract (**Figure 7A**). pTh17 were in close localization with NLRP3+ AREG+ MNPs in PCD fistula tracts (**Figure 7B**). Notably, these cells were present in all PCD samples but were scarce in IPF samples. Importantly, ligand-receptor analysis revealed elevated IFN-γ signaling in PCD spatial niches populated by pTh17 and NLRP3+ AREG+ MNPs (**Figure 7C**). Taken together, our ST data demonstrate PCD-specific spatial niches harboring both pTh17s and NLRP3+ AREG+ MNPs with elevated IFN-γ signaling.

**Figure 7.**
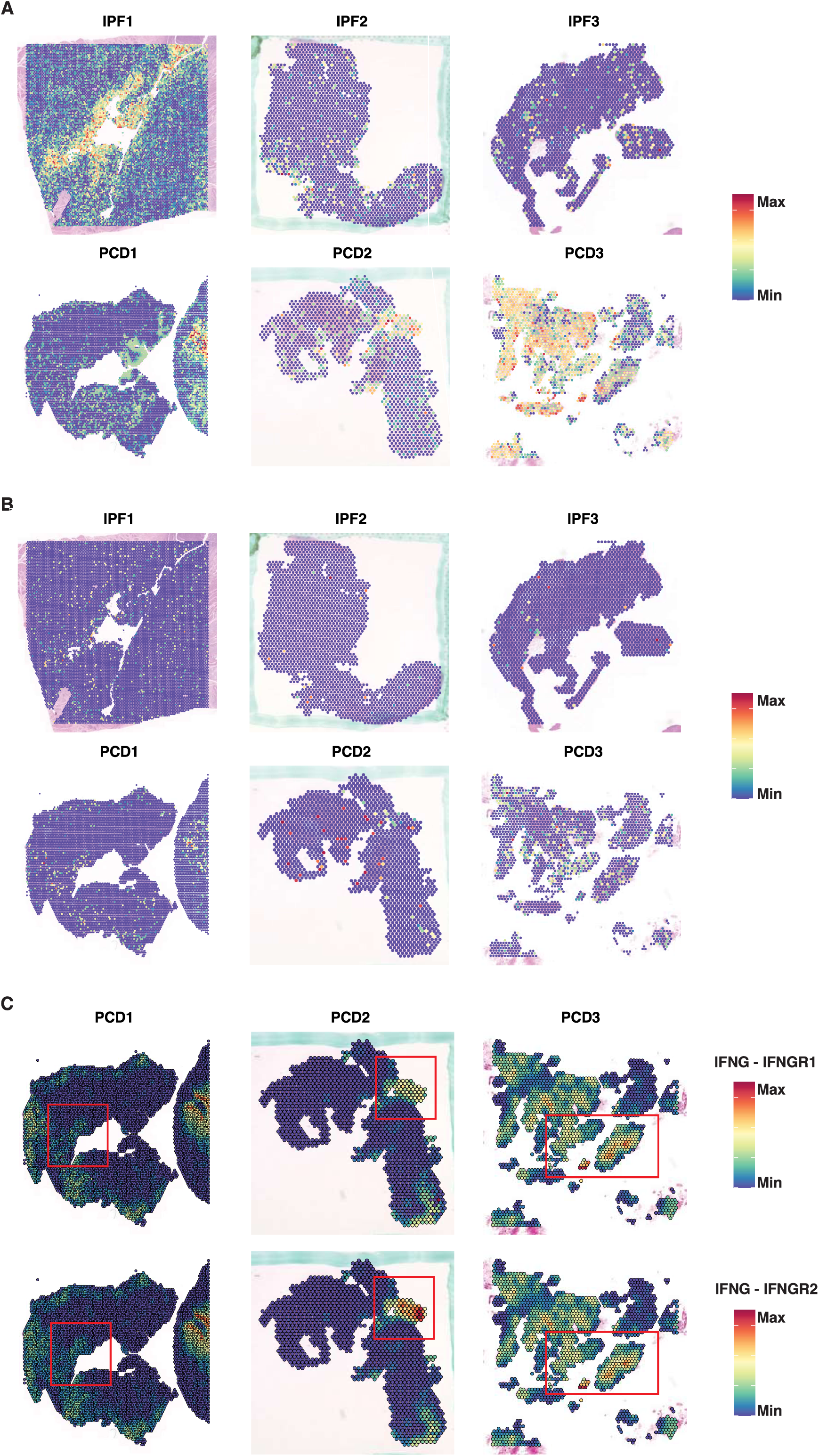
NLRP3+ AREG+ MNPs and pTh17 spatially co-localize and drive local IFN-γ signaling. (A) Single-spot score of **NLRP3+ AREG+** MNP gene module in IPF and PCD fistulas. (B) Single-spot score of pTh17 gene module in IPF and PCD fistula. (C) Spatial activity of IFNG-IFNGR1 and IFNG-IFNGR2 signaling pairs.

## Discussion

PCD is a refractory presentation of CD where current therapies are mainly approved through registrational trials in luminal CD and then subsequently used in the treatment of patients with PCD. This is primarily due to limited knowledge about the pathophysiology of PCD unlike luminal CD. Given this unmet need, we defined the mucosal single-cell landscapes of PCD in an extensive patient cohort using scRNA-seq, CyTOF, spatial transcriptomics, and IHC.

ScRNA-seq of fistula tissues revealed pathogenic pathways, especially elevated IFN-γ and TNF-α signaling in the lymphocyte and myeloid cell compartments in PCD. In addition to fistula tracts, we showed enhanced IFN-γ and TNF-α pathway activities in the ileal mucosa of PCD patients compared to patients with NPCD, and in rectal mucosa associated with active PCD vs. inactive/healed PCD. Although TNF-α activation has long been linked to PCD^23^, the evidence of IFN-γ involvement was lacking. Prior studies have also suggested that genetic contributions to the CD fistulizing phenotype differ in perianal and internal fistulae. Given augmented IFN-γ expression and response shared features in both fistula tracts and luminal mucosa in our study, it is important to determine whether there are common factors that predispose to fistulization in internal penetrating and perianal fistulas. Future studies should also identify the underlying mechanisms, including host factors (e.g., genetic and epigenetic changes), environmental stimuli (e.g., smoking, air pollutants), microorganisms (e.g., skin microflora), and medications (e.g., prior and current use of immunosuppressants), which trigger the elevations in IFN-γ secretion and/or responsiveness in PCD.

EMT is implicated in PCD pathogenesis, although the molecular and cellular mechanisms remain limited^3^. Here, we identified active PCD-enriched epithelial cell populations in the rectum, which overexpressed IFN-γ/TNF-α-receptors and responsive genes such as IFNGR1/2, TNFRSF1A, STAT1, and NFKB1/2, as well as EMT genes that were transcriptionally activated by IFN-γ/TNF-α signaling. Our data thus suggest a direct upregulation of EMT by IFN-γ and TNF-α in PCD rectum. ST further found that fistula epithelial cells corresponded to IFN-γ- and TNF-α-responsive rectal epithelial cells, further supporting the pathogenic role of epithelial IFN-γ and TNF signaling in perianal fistulization. Ongoing studies are investigating how IFN-γ and TNF-α work in synergy to promote EMT in rectal and fistula epithelial cells from PCD patients using an organoid culture model.

In PCD patients, IFN-γ-producing pTh17 cells driven by cytokine and antigen stimulation may play a central role in the initiation, progression, and maintenance of perianal fistulas. By analyzing the APCs in our dataset, we identified elevated NLRP3 and AREG in TREM1+ CD172a+ myeloid cells in PCD, which may represent macrophages associated with therapeutic resistance in CD patients in the literature^30, 53–57^. Increased single-cell scores for LPS signaling in NLRP3+ AREG+ cells and myeloid cells as a whole indicate an altered anti-microbial response in the development of pTh17 cells and PCD pathogenesis. Ligand-receptor analysis also revealed an upregulation of APC-derived mediators that promote the differentiation (e.g., IL6, IL1B), activation (e.g., IL15)^34^, and recruitment (e.g., CCL2/8, CXCL9/10)^31–33^ of Th17 cells in PCD fistulas. Furthermore, spatial analysis demonstrated a close localization of pTh17 with NLRP3+ AREG+ MNPs, which further correlates with elevated IFNG signaling in a PCD-specific manner.

Using CyTOF, we corroborated our scRNA-seq findings and demonstrated additional features of TNF and IFN-γ signaling in PCD. The fistula tract, external fistula opening, and rectum in PCD patients exhibited enrichment of CD172a/b+ TREM1+ MNPs, which were LPS-responsive cells by scRNA-seq and ST and responsible for chronic antigen stimulation of Th17 cells. Interestingly, circulating Th17 cells is only elevated in the fistula tracts in PCD, as are markers of IFN-γ and TNF-α production and T cell activation. This suggests a mechanism where LPS-responsive APCs recruit Th17 cells and contribute to sustenance of the fistula tract.

The hyperactivation of IFNG expression and response suggest that blocking IFNG pathways is a promising therapeutic target for PCD. Indeed, a new Janus kinase (JAK) inhibitor, upadacitinib, has shown efficacy in patients with PCD in clinical trials and real-world practice^58,59^. The JAK-STATs are essential components of IFN-γ response in immune, stromal, and epithelial cells^60^; it is possible that upadacitinib induces perianal fistula healing by suppressing IFN-γ pathways. However, even with upadacitinib, the rates of perianal fistula healing were low even in the 30 mg maintenance dose arm, with resolution of drainage seen in 23.1% and closure of perianal fistula external openings only in 16.0%^59^. A humanized monoclonal antibody for IFN-γ, fontolizumab, was previously tested on patients with CD^61–63^. Treatment with fontolizumab was well tolerated with significant reductions in C-reactive protein (an inflammatory marker), although a strong clinical response was not observed^63^. It is worth noting that no data regarding the presence of perianal disease was available in the last trial. Therefore, with the low rates of fistula healing with current therapies, clinical trials of systemic IFN-γ antagonists for PCD patients are warranted.

Our study also identified two gene expression modules associated with use of anti-TNFs at the fistula level through a scRNA-seq approach. While a prior study has noted the absence of anti-TNF drugs in perianal fistula tissue in patients non-responsive to anti-TNFs, no previous studies specifically examined local tissue responses to anti-TNFs^64^. Further prospective sample recruitment is ongoing to validate the findings and extend to other mechanisms of actions.

The strengths of our study include the presence of two control groups, the IPF and the NPCD groups across multiple methodologies for single-cell characterization of PCD, contrasted against IPF and NPCD. Furthermore, we compared our findings against known and published cohorts of scRNA-seq and ST in CD including those with and without perianal disease. Additionally, active disease activity in the rectum has been associated with lower rates of fistula healing^65^. This is supported by our findings that inflamed rectal mucosa exhibited hyperactivated IFN-γ and TNF-α responses and EMT. Limitations of our study include the likelihood that the biopsies obtained from the fistula tract may not be representative of the entire fistula tract. However, we have attempted to compare findings from within the fistula tract to the external opening of the tract in a subset of the PCD patients. Additionally, duration of disease may influence epithelization of the fistula tracts. However, the mean disease duration was similar across the 3 groups studied. Finally, it remains unclear how therapy impact the study findings; however we have presented analyses comparing those on anti-TNFs and advanced therapy naïve in a subset of PCD patients.

## Conclusion

In this study, we performed scRNA-seq, CyTOF, spatial transcriptomics, and IHC for a comprehensive and detailed analysis of PCD, a challenging condition with unclear pathophysiology to date. PCD fistulas showed elevated pathogenic pathways including IFN-γ and TNF-α responses in myeloid and stromal cells. In addition to fistula tracts, intestinal cells from PCD patients also expressed greater levels of IFN-γ-responsive and EMT genes compared to NPCD. In addition, we showed that both fistula tracts and ileal mucosa from PCD patients harbored expanded IFNG+ pathogenic Th17 cells, which expressed augmented inflammatory cytokines and chemokines. Skewed immune responses including increased Th17 cells, elevated proinflammatory myeloid cells, and altered T cell exhaustion markers in the fistula tracts and surrounding tissues in PCD patients were identified by CyTOF. Further analysis revealed cellular modules associated with anti-TNF therapy in PCD patients. Our findings suggest a therapeutic potential of suppressing IFN-γ and related pathways in inducing and maintaining remission in PCD patients.

## Acknowledgements

The work was made possible by the generous supports of the American Gastroenterological Association Fellowship-to-Faculty Transition Award (AGA2023-32-03, Siyan Cao), Digestive Disease Research Core Center (DDRCC) Pilot and Feasibility Award (P30 DK052574, Siyan Cao), NIH/NIDDK K08 Clinical Investigator Award (1K08DK140612-01, Siyan Cao), and Washington University (WU) DDRCC (NIDDK P30 DK052574). The authors thank the stellar clinical research coordinators: Darren Billy Nix, Donald Jones, Ginny Van Teslaar, Vicki Martin, and Guadalupe Oliva Escudero at WU Inflammatory Bowel Disease Center for help with IRB, recruitment of patients, and sample collection. We thank the WU Genome Access Technology Center and Immunomonitoring Laboratory for help with scRNA-seq/spatial transcriptomics and CyTOF, respectively. Siyan Cao was also supported by a Crohn’s and Colitis Foundation Career Development Award (CCF1062472), WU Clinical and Translational Research Funding Program (CTRFP1714), WU Precision Health Innovation Award (PHIA 101), Lawrence C. Pakula, MD IBD Education & Innovation Fund, and Doris Duke COVID-19 Fund to Retain Clinical Scientists Program. Parakkal Deepak was supported by a Junior Faculty Development Award from the American College of Gastroenterology and IBD Plexus of the Crohn’s and Colitis Foundation. Marco Colonna was supported by NIH R01 DK126969 and R01 DK132327. Matthew A. Ciorba was supported by Givin’ It All For Guts Foundation. Ta-Chiang Liu was supported by NIH R01 DK125296, DK124274, DK136829, and DK138465.

## Author Contributions

**Conceptualization:** SC, PD.

**Data curation:** SC, KN (equal).

**Formal analysis:** KN, SC (equal).

**Funding acquisition:** SC, PD.

**Investigation:** KN, SC (equal).

**Methodology:** SC, PD, KN (equal).

**Project administration:** SC, PD.

**Resources:** SC, KM, PD, GOE, DMJ, VLM, GVT, DN, RS, MS, SRH, PEW, MGM, SCG, WCC, MC, and PD.

**Software:** KN.

**Supervision:** SC, PD.

**Visualization:** YL, KN, SC.

**Writing – original draft:** KN, SC, PD (equal).

**Writing – review and editing:** KN, SC, PD, MC (equal).

## Data Transparency Statement

We will make our data, analytic methods, and study materials available to other researchers. The Transcript profiling data has been submitted to NCBI’s GEO (GSE277387).

