## Supplemental methods for "Single-Cell and Spatial Multi-omics Reveal Interferon Signaling in the Pathogenesis of Perianal Fistulizing Crohn’s Disease"

### Sample cryopreservation

Briefly, the biopsies were kept in complete RPMI medium on ice immediately after collection and transported to the lab, where they were changed into a freezing medium (10% dimethyl sulfoxide in fetal calf serum), transferred to a pre-chilled Mr. Frosty™ (ThermoFisher) container with isopropanol, and kept in -80°C. The samples were then transferred to liquid nitrogen 24h later for storage. The cryopreservation of samples, isolation of immune cells, and staining for CyTOF followed the sample protocols we recently described[10]. Briefly, the biopsies were kept in complete RPMI medium on ice after collection and immediately transported to the lab, where they were changed into a freezing medium with 10% dimethyl sulfoxide, frozen in -80°C in a freezing container and transferred to liquid nitrogen 24h later. To minimize variation, the samples were processed in batches as described before[10]. Briefly, the cryopreserved specimens were gently thawed in 37°C water bath, mucosal immune cells were extracted by digesting tissues in complete RPMI medium containing collagenase IV at 37°C for 50 min under agitation. Cells were then filtered and subjected to density gradient centrifugation using Percoll solutions.

### CyTOF analysis

Samples were manually gated using Cytobank as described to exclude background, dead cells (Cisplatin+), doublets (DNA1/2 stain), and normalization beads. Dimensionality reduction was performed on CD45+ CD3+ CD19- T cells, CD45+ CD3- CD19+ B cells, and CD45+ CD3- CD19- innate immune cells to identify cell clusters based on surface markers, using the viSNE tool in Cytobank (**Supplementary Figure 4A**). Circulating Th17s were defined as CD103- CD45RO+ CD161+ CD127+ CD4+ T cells, as in [1]. CD172a/b+ TREM1+ MNPs were defined as CD14+ CD33+ CD172a/b+ TREM1+ innate immune cells. Tc17 cells were defined as CD26+

CD161+ CD8+ T cells. Cell populations expressing certain markers (e.g. CD39+ CD4+) were calculated by summing counts of clusters expressing these proteins.

**Library preparation and sequencing**

scRNA-seq, TCR-seq, and BCR-seq library was prepared using the 10X Chromium Next GEM Single Cell 5' Kit v2, Chromium Single Cell Human TCR Amplification Kit, and Chromium Single Cell Human BCR Amplification Kit, respectively. Spatial transcriptomic library was prepared using the 10X Visium V4 – FFPE V2 chemistry. Sequencing was performed on a NovaSeq X Plus sequencer.

**scRNA-seq data preprocessing, quality control, and cluster annotation**

scRNA-seq data was processed using the CellRanger suite (v7.1.1), and count matrices were loaded into Seurat (v4.4.0)[2]. Cells with < 1000 or > 4000 unique detected features, > 30000 transcripts, and > 10% percent mitochondrial genes were discarded. Doublets were removed using DoubletFinder[3]. Normalization was done using SCTransform[4], and batch correction was performed using the reciprocal PCA approach in Seurat. Differential expression testing was done using the Wilcoxon Rank Sum test.

Following dimensionality reduction and initial clustering, coarse cell type clusters were identified as follows: B cells (MS4A1), plasma cells (SDC1), T cells (CD3D), ILCs (CD3D- cells that clustered closely to T cells), mononuclear phagocytes (ITGAX), plasmacytoid dendritic cells (pDCs; CLEC4C), mast cells (KIT, TPSAB1, TPSB2), endothelial cells (VWF), stromal cells (TAGLN). Subclustering analysis was performed by re-clustering B and plasma cells, T cells and ILCs, myeloid cells (mononuclear phagocytes, pDCs and mast cells), and stromal cells. Subclusters were annotated as described in the main text. A complete list of marker genes for each cluster is included in **Supplementary Table 4.**

**Spatial transcriptomics (ST) analysis**

Spatial transcriptomics data was analyzed using Seurat (v4.4.0). Count data from individual samples were loaded and normalized using SCTransform[4]. Normalized data was then merged into a single Seurat object and integrated using the canonical correlation analysis (CCA) method in Seurat. Cluster marker genes were identified using the FindConservedMarker function and the Wilcoxon Rank Sum test.

**Reanalysis of published datasets from Washburn et al.[5] and Kong et al.[5]**

Washburn et al. dataset: Data was loaded into Seurat, and quality control was performed as described by the authors. Data was normalized using log transformation and batch-corrected using Harmony[6]. Ambient RNA contamination was corrected using SoupX[7], and technical dropouts were imputed using adaptively thresholded low-rank approximation[8]. Non-epithelial cell clusters were annotated using Seurat's label transfer, with our in-house dataset as reference. Epithelial cells were identified based on KRT8 expression. Epithelial cell subclusters (as in Supplementary Fig 3) were annotated as follows: Colonocytes (C0, 2), IFNGR+ TNFR+ colonocytes (C1), cycling (C3, 7, 8), Goblet cells (C4, 5), crypt cells (C6), IFN-responsive (C9), BEST4+ epithelial cells (C10), TNF-responsive (C12).

Kong et al. dataset: Data was loaded into Seurat and quality control was performed as described by the authors. Data was normalized using log transformation and batch-corrected with Harmony.

**Pathway analysis, module scoring, and transcription factory activity inference.**

Pathway analysis was performed via the SCPA R package[9], using the Hallmark human gene sets. Single-cell module scores were calculated using the Mann-Whitney U statistic[10]. Interferon gamma signaling, TNF-a signaling, and epithelial-mesenchymal transition scores (as in Figure 3) were calculated using the GOBP_RESPONSE_TO_TYPE_II_INTERFERON, GOBP_TUMOR_NECROSIS_FACTOR_MEDIATED_SIGNALING_PATHWAY, and

GOBP_EPITHELIAL_TO_MESENCHYMAL_TRANSITION gene sets from MsigDB[11], respectively. Pathogenic Th17 (pTh17) gene module (as in Figure 7) consisted of CD3D, IL17A, and IFNG, while the NLRP3+ AREG+ MNP gene module consisted of CD14, FCGR3A (encoding CD16), NLRP3, AREG, and TREM1. Transcription factor activity was inferred using the Python implementation of SCENIC (pySCENIC[12]).

**Ligand-receptor signaling analysis**

Ligand-receptor (LR) signaling analysis was performed using NICHES[13]. Prior to analysis, normalized and annotated scRNA-seq/ST data underwent imputation using adaptively thresholded low-rank approximation[8] to correct for technical dropouts. NICHES was ran using the fantom5 ligand-receptor database. Cluster marker pathways were identified using the receiver operating characteristic (ROC) test within Seurat's FindMarkers function, using an AUC threshold of 0.8.

**Weighted gene co-expression network analysis**

IPF samples were excluded from the scRNA-seq dataset, as was one PCD sample from a patient receiving Ustekinumab, resulting in a total of 8 samples (4/group). Raw scRNA-seq counts were pseudobulked using the summation method, and genes that were unannotated or encoding lncRNA were excluded. Pseudobulked matrix was then batch-corrected using ComBat_seq[14] with anti-TNF treatment status as a covariate. Adequate batch correction was confirmed using PCA. The matrix was then normalized using the regularized log transformation method in DESeq2[15], and genes with low variance were removed. The adjacency matrix was constructed using softPower of 9. Following construction of module eigengenes, modules with less than 25% dissimilarity were merged, resulting in a total of 32 modules.

**Cell type deconvolution for ST**

Spot deconvolution from ST data was performed using CARD[16]. Genes with under 100 counts and spots with under 5 counts were excluded from deconvolution, and deconvolution was performed on each spatial sample separately. The single-cell reference was generated by combining our fistula scRNA-seq data and Washburn et al[5] rectal scRNA-seq dataset, to account for the lack of epithelial cells in our dataset. Briefly, the rectal dataset underwent SCTransform for consistency with our in-house data; the datasets were then integrated in Seurat using reciprocal PCA. A simplified cell annotation was used: Tfh, Treg, Th17, and naïve CD4/Tcm were grouped together as CD4 T cells; CD8 Trm precursor, CD8 Trm, CD8 Tem, MAIT were grouped together as CD8 T cells; CD56dim NK, CD56hi NK, ILC3 were grouped together as ILCs; naïve and memory B cells were grouped together as B cells; cDC1/2 and mRegDCs were grouped together as DCs; and inflammatory-fibroblast-like, PDGFRAhi fibroblasts, and DPT+PI16+ fibroblasts were grouped together as fibroblasts. Raw counts were inputted for CARD deconvolution.

6. Korsunsky I, Millard N, Fan J, et al. Fast, sensitive and accurate integration of single-cell data with Harmony. Nature methods 2019;16:1289-1296.
7. Young MD, Behjati S. SoupX removes ambient RNA contamination from droplet-based single-cell RNA sequencing data. Gigascience 2020;9:giaa151.
8. Linderman GC, Zhao J, Roulis M, et al. Zero-preserving imputation of single-cell RNA-seq data. Nature communications 2022;13:192.
9. Bibby JA, Agarwal D, Freiwald T, et al. Systematic single-cell pathway analysis to characterize early T cell activation. Cell Reports 2022;41.
10. Andreatta M, Carmona SJ. UCell: Robust and scalable single-cell gene signature scoring. Computational and structural biotechnology journal 2021;19:3796-3798.
11. Liberzon A, Subramanian A, Pinchback R, et al. Molecular signatures database (MSigDB) 3.0. Bioinformatics 2011;27:1739-1740.
12. Van de Sande B, Flerin C, Davie K, et al. A scalable SCENIC workflow for single-cell gene regulatory network analysis. Nature protocols 2020;15:2247-2276.
13. Raredon MSB, Yang J, Kothapalli N, et al. Comprehensive visualization of cell–cell interactions in single-cell and spatial transcriptomics with NICHES. Bioinformatics 2023;39:btac775.
14. Zhang Y, Parmigiani G, Johnson WE. ComBat-seq: batch effect adjustment for RNA-seq count data. NAR genomics and bioinformatics 2020;2:lqaa078.
15. Love MI, Huber W, Anders S. Moderated estimation of fold change and dispersion for RNA-seq data with DESeq2. Genome biology 2014;15:1-21.
16. Ma Y, Zhou X. Spatially informed cell-type deconvolution for spatial transcriptomics. Nature biotechnology 2022;40:1349-1359.
