## Supplemental figure 1 for "Single-Cell and Spatial Multi-omics Reveal Interferon Signaling in the Pathogenesis of Perianal Fistulizing Crohn’s Disease"

A

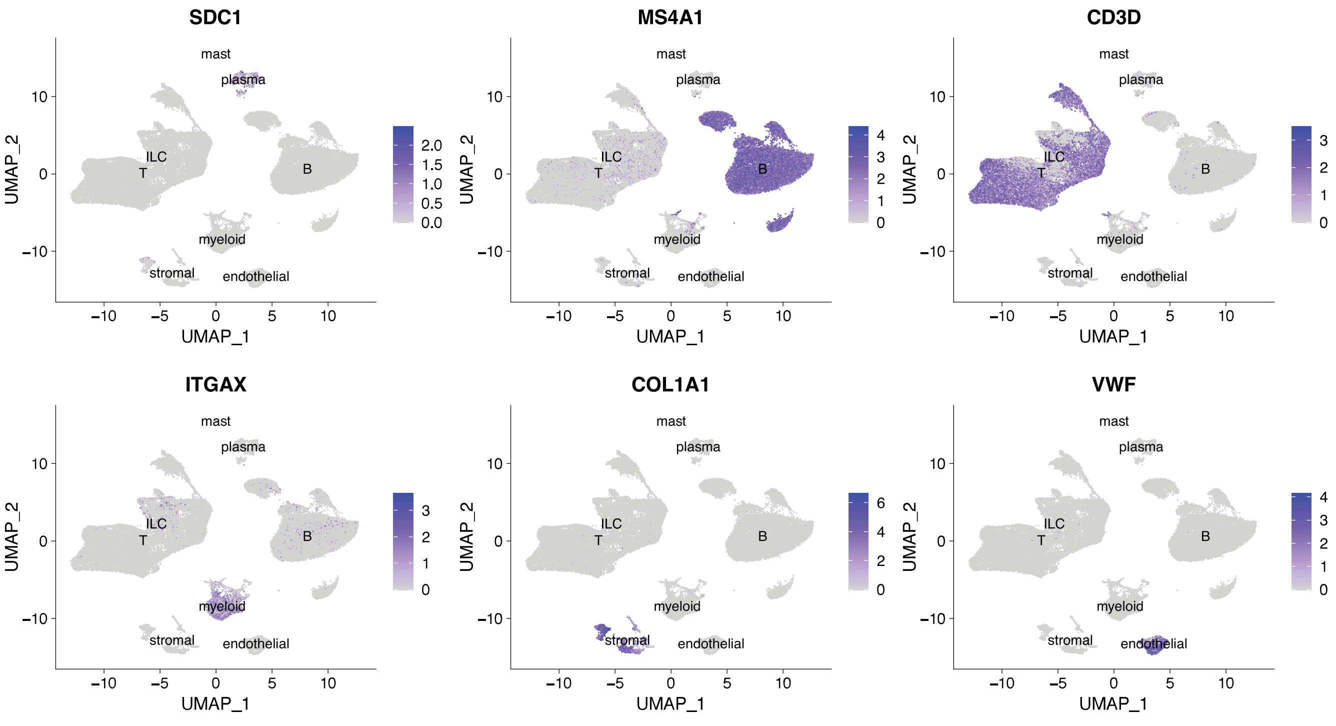


**Supplementary Figure 1 (related to Figure 1).** UMAP plots indicating expression of canonical markers used to identify the major cell compartments in the fistula tract.
