## Supplemental figure 2 for "Single-Cell and Spatial Multi-omics Reveal Interferon Signaling in the Pathogenesis of Perianal Fistulizing Crohn’s Disease"

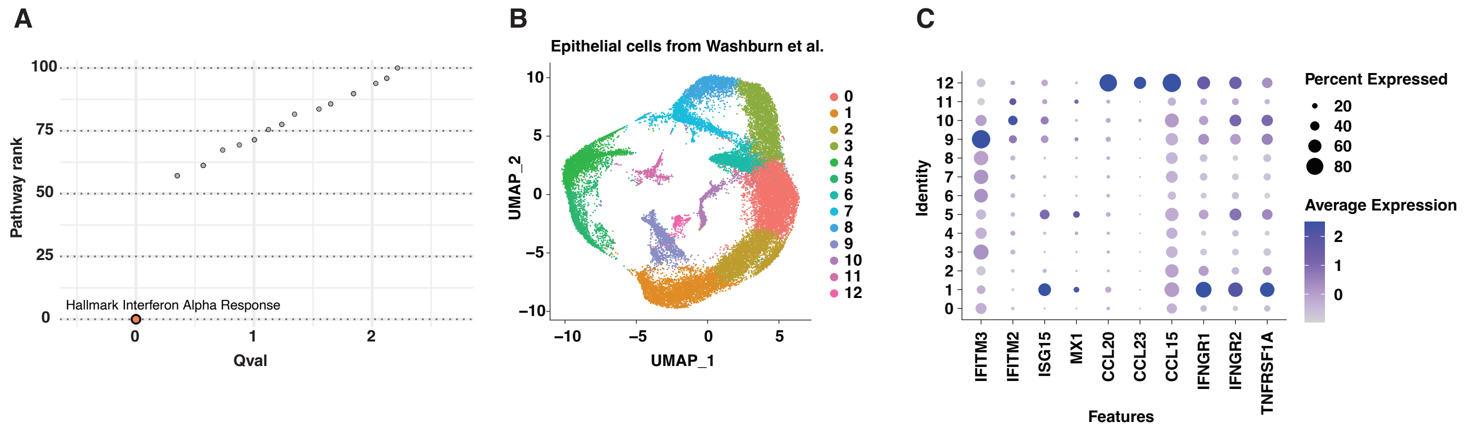


**Supplementary Figure 2 (related to Figure 2).** (A) Pathway rank of IFNA response in PCD fistulas. (B) UMAP plot of rectal epithelial cells from Washburn et al. (C) Expression of interferon response genes of rectal epithelial cell clusters in (B).
