## Supplemental figure 3 for "Single-Cell and Spatial Multi-omics Reveal Interferon Signaling in the Pathogenesis of Perianal Fistulizing Crohn’s Disease"

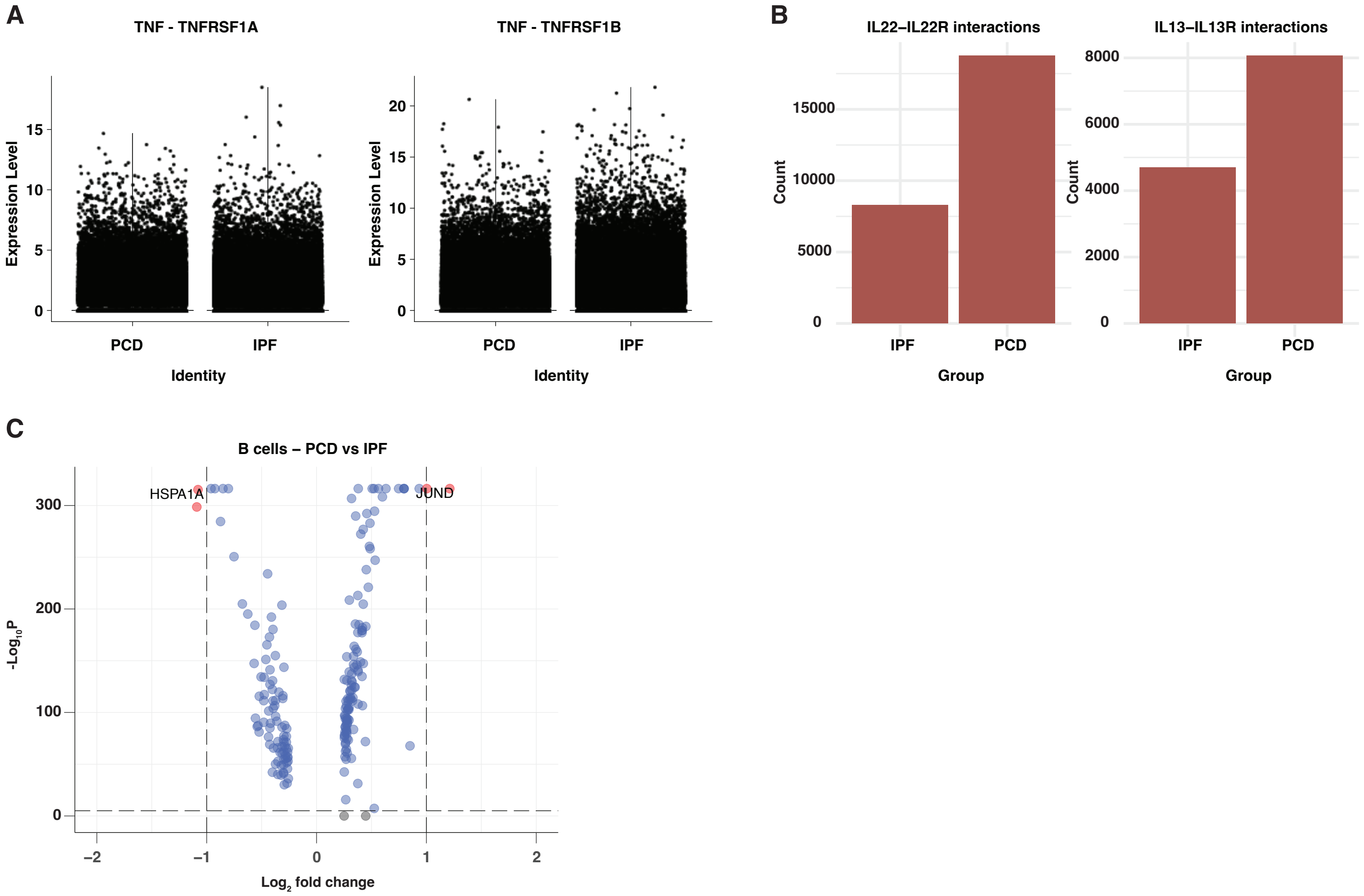


**Supplementary Figure 3 (related to Figure 3).** (A) TNF signaling in PCD and IPF fistula tracts. (B) Total counts of IL22-IL22R and IL13-IL13R interacting pairs in PCD vs IPF. (C) Volcano plot of DEGs in B cells in PCD vs. IPF.
