## Supplemental figure 4 for "Single-Cell and Spatial Multi-omics Reveal Interferon Signaling in the Pathogenesis of Perianal Fistulizing Crohn’s Disease"

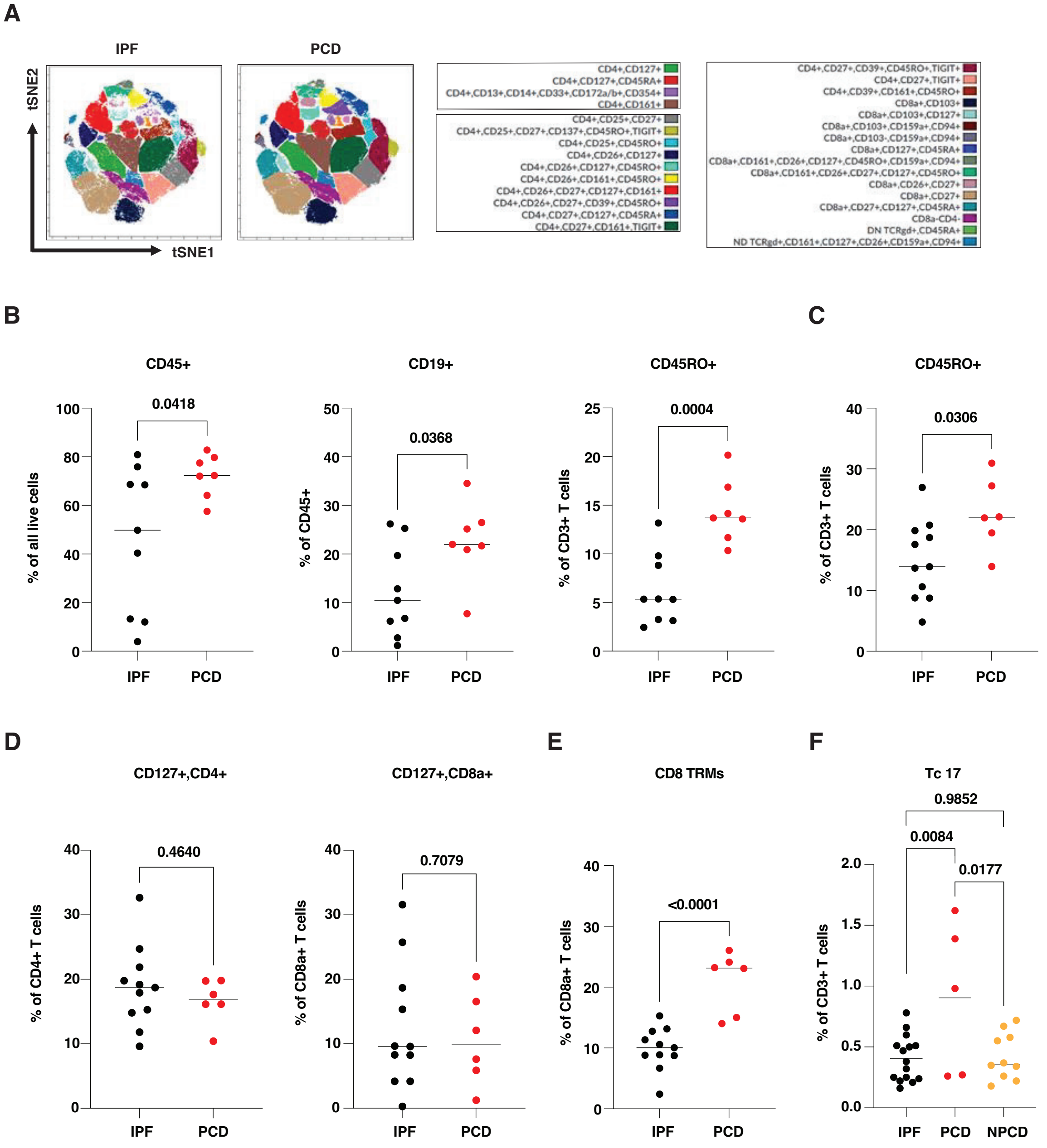


**Supplementary Figure 4 (related to Figure 4).** (A) Representative tSNE plot of CyTOF cell clusters identified using viSNE. (B-E) Mucosal immune cells populations analyzed by CyTOF. (B) Fistula tracts. (C-E) External fistula opening. (F) Rectum.
