## Supplemental figure 5 for "Single-Cell and Spatial Multi-omics Reveal Interferon Signaling in the Pathogenesis of Perianal Fistulizing Crohn’s Disease"

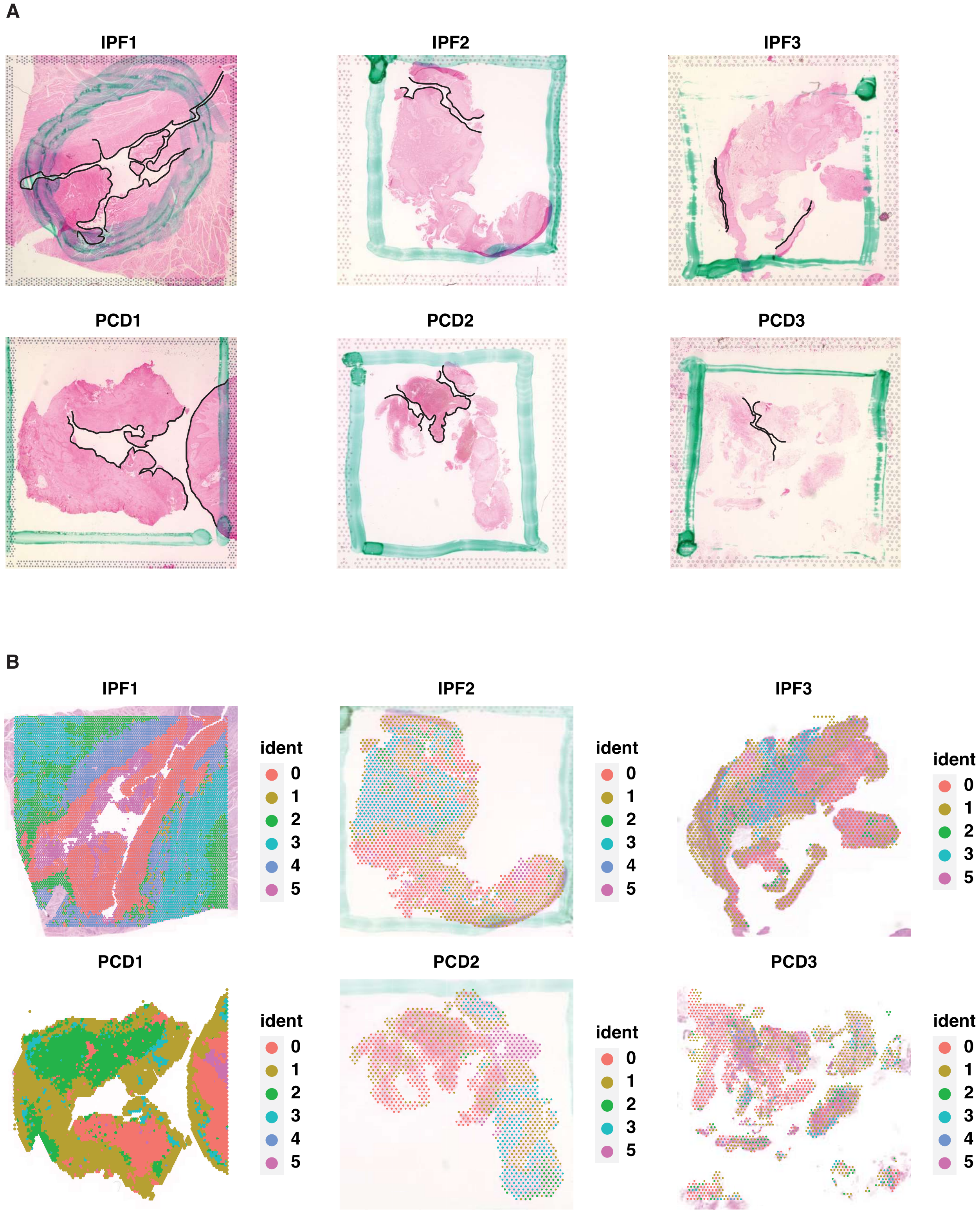


**Supplementary Figure 5 (related to Figure 6).** (A) H&E staining of IPF and PCD fistula samples analyzed using ST; black lines denote fistula tract lining. (B) Spatial plot of clusters in IPF and PCD fistula as identified by ST.
