## Supplemental figure 6 for "Single-Cell and Spatial Multi-omics Reveal Interferon Signaling in the Pathogenesis of Perianal Fistulizing Crohn’s Disease"

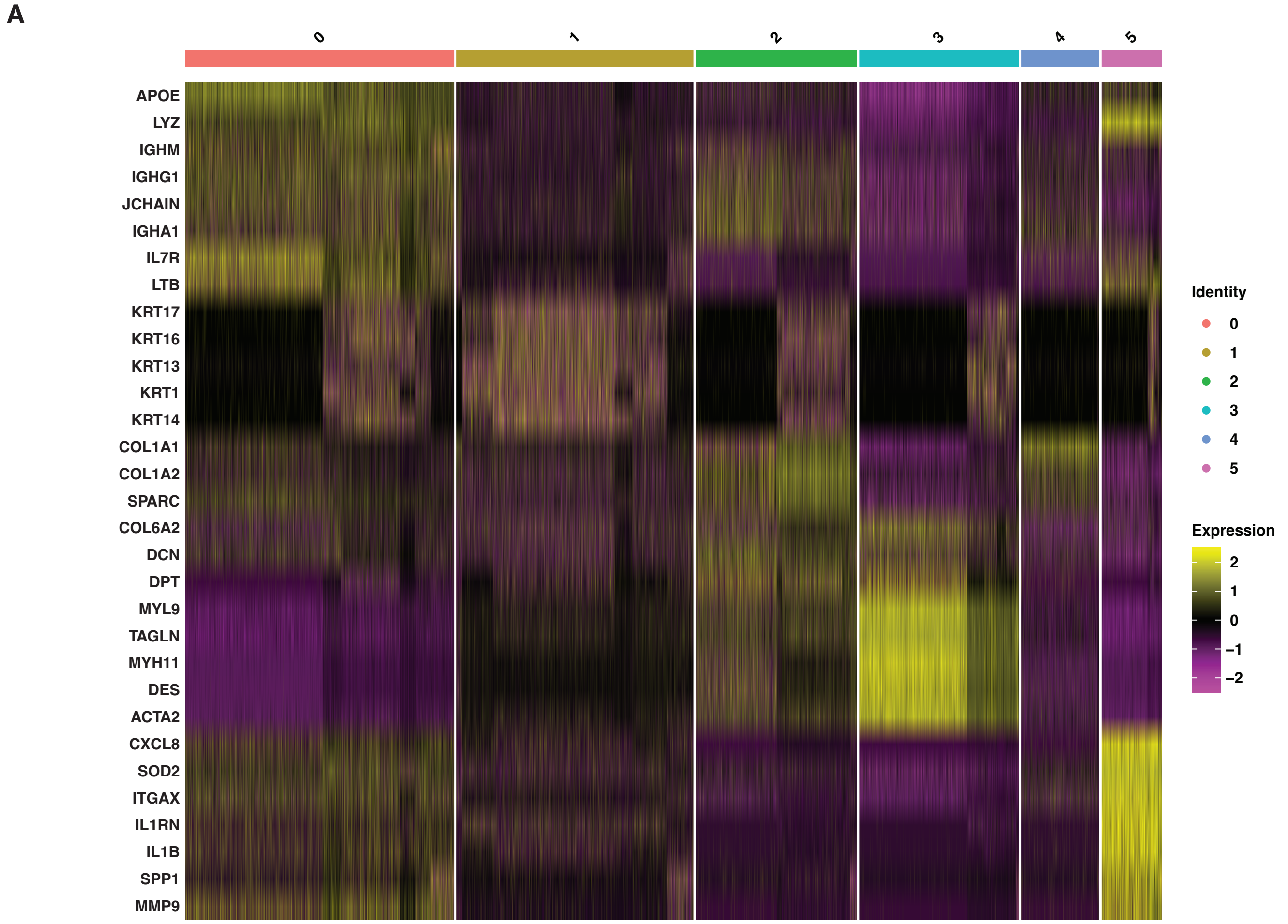


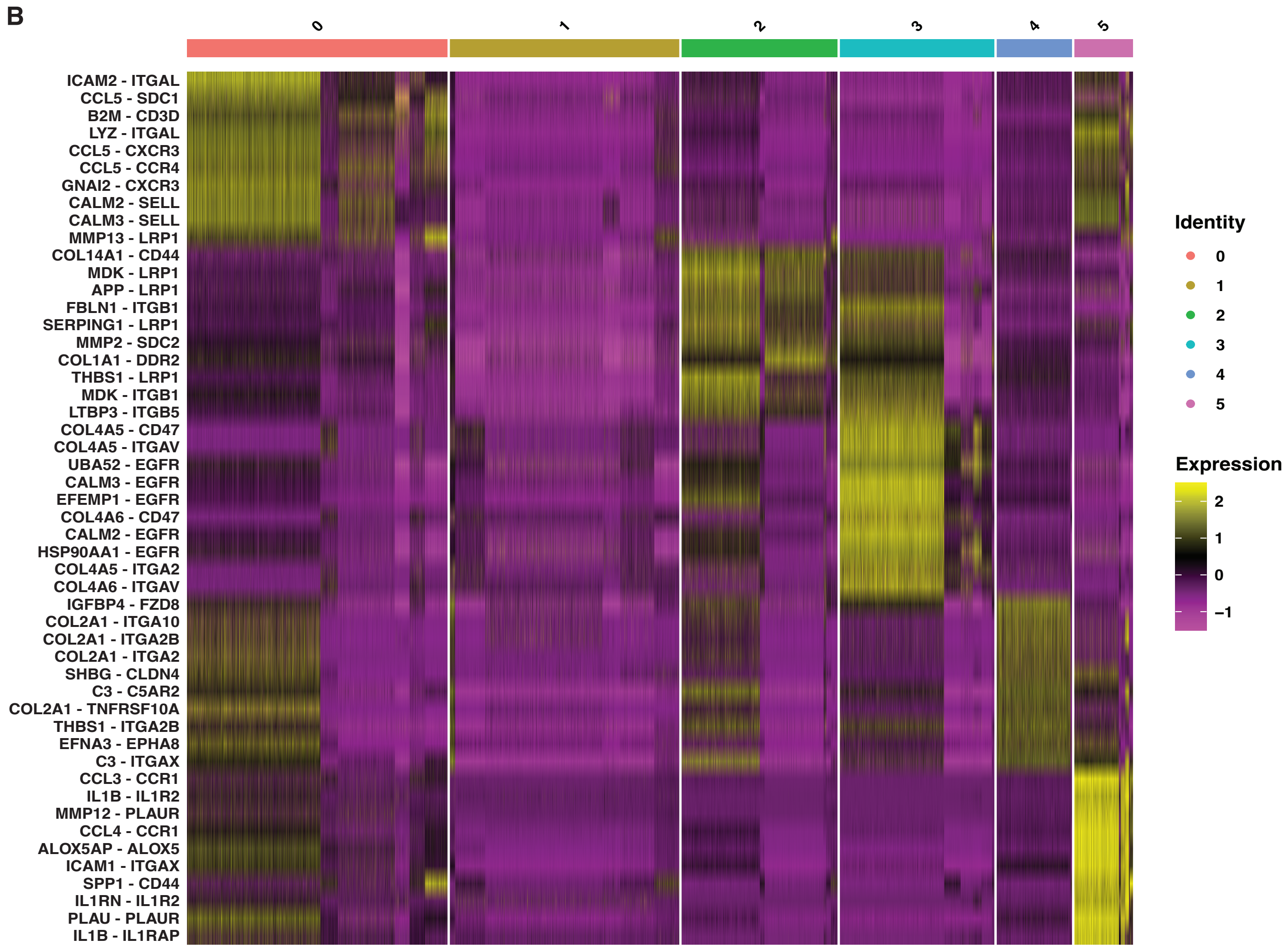


**Supplementary Figure 6 (related to Figure 6).** (A) Heatmap of marker gene expression across spatial clusters. (B) Heatmap of marker ligand-receptor signaling pairs across spatial clusters.
