## Supplemental table 1 for "Single-Cell and Spatial Multi-omics Reveal Interferon Signaling in the Pathogenesis of Perianal Fistulizing Crohn’s Disease"

| | Perianal fistulizing CD (N=24) | CD without perianal disease (N=10) | Idiopathic/cryptoglandular fistula (N=29) |
|---|---|---|---|
| **Age, mean (SD), y** | 40.9 (11.0) | 43.3 (11.1) | 47.0 (14.6) |
| **Sex, N (%)** | | | |
| Female | 14 (58.3) | 4 (40.0) | 10 (34.5) |
| Male | 10 (41.7) | 6 (60.0) | 19 (65.5) |
| **Race, N (%)** | | | |
| Asian | 1 (4.2) | 0 (0.0) | 1 (3.4) |
| Black or African American | 3 (12.5) | 1 (10.0) | 3 (10.3) |
| Other Pacific Islander | 0 (0.0) | 0 (0.0) | 0 (0.0) |
| White | 20 (83.3) | 9 (90.0) | 25 (86.2) |
| **Tobacco Use, N (%)** | | | |
| Current | 7 (29.2) | 1 (10.0) | 6 (20.7) |
| Former | 4 (16.7) | 4 (40.0) | 7 (24.1) |
| Never | 13 (54.2) | 5 (50.0) | 16 (55.2) |
| **Duration of Diagnosis, mean (SD), y** | 16.2 (11.2) | 16.3 (9.4) | 4.6 (8.2) |
| **Age at diagnosis, N (%)** | | | |
| A1 [<16 y] | 3 (12.5) | 2 (20.0) | 0 (0.0) |
| A2 [17-40 y] | 20 (83.3) | 8 (80.0) | 15 (51.7) |
| A3 [>40 y] | 1 (4.2) | 0 (0.0) | 14 (48.3) |
| **Location, N (%)** | | | |
| L1 | 0 (0.0) | 1 (10.0) | 0 (0.0) |
| L2 | 7 (29.2) | 1 (10.0) | 0 (0.0) |
| L3 | 17 (70.8) | 7 (70.0) | 0 (0.0) |
| L4 | 0 (0.0) | 1 (10.0) | 0 (0.0) |
| None | 0 (0.0) | 0 (0.0) | 29 (100.0) |
| **Behavior, N (%)** | | | |
| B1 | 8 (33.3) | 3 (30.0) | 0 (0.0) |
| B2 | 6 (25.0) | 4 (40.0) | 0 (0.0) |
| B3 | 10 (41.7) | 3 (30.0) | 0 (0.0) |
| None | 0 (0.0) | 0 (0.0) | 29 (100.0) |
| **Antidiarrheal, N (%)** | | | |
| Yes | 11 (45.8) | 3 (30.0) | 5 (17.2) |
| No | 13 (54.2) | 7 (70.0) | 24 (82.8) |
| **5-Aminosalicylic Acid, N (%)** | | | |
| Yes | 6 (25.0) | 3 (30.0) | 0 (0.0) |
| No | 18 (75.0) | 7 (70.0) | 29 (100.0) |
| **Biologics, N (%)** | | | |
| Adalimumab | 3 (12.0) | 1 (10.0) | 0 (0.0) |
| Infliximab | 4 (16.0)* | 1 (10.0) | 0 (0.0) |
| Ustekinumab | 6 (24.0) | 3 (30.0) | 0 (0.0) |
| Vedolizumab | 1 (4.0) | 1 (10.0) | 0 (0.0) |
| None | 11 (44.0) | 4 (40.0) | 29 (100.0) |
| **Corticosteroids** | | | |
| Yes | 3 (12.5) | 3 (30.0) | 1 (3.4) |
| No | 21 (87.5) | 7 (70.0) | 28 (96.6) |
| **Immunomodulators** | | | |
| Yes | 4 (16.7) | 6 (60.0) | 0 (0.0) |
| No | 20 (83.3) | 4 (40.0) | 29 (100.0) |
| **Antibiotics** | | | |
| Yes | 4 (16.7) | 6 (60.0) | 7 (24.1) |
| No | 20 (83.3) | 4 (40.0) | 22 (75.9) |
| **Proctitis Endoscopy** | 5 (20.8) | 2 (20.0) | 0 (0.0) |
| **Proctitis Histology** | 6 (25.0) | 1 (10.0) | 0 (0.0) |
| **Proctitis MRI** | 3 (12.5) | 1 (10.0) | 1 (3.4) |
| **Sample Type$^{+}$** | | | |
| CyTOF | 7 (28.0) | 10 (100.0) | 15 (50.0) |
| scRNA-seq | 9 (36.0) | 0 (0.0) | 6 (20.0) |
| Spatial transcriptomics | 3 (12.0) | 0 (0.0) | 3 (10.0) |
| IHS | 6 (24.0) | 0 (0.0) | 6 (20.0) |

**Supplemental Table 1: Clinical characteristics of patients.**
*One patient with two samples at different clinical times was on infliximab at the time of one of the biopsies and was not on any biologics at the second biopsy.

+There were 25 samples for perianal Crohn's disease for a total of 24 patients, and 30 samples were analyzed for 29 patients in the cryptoglandular fistula group (one patient in each group had two samples).
